# Epileptic seizure suppression: a computational approach for identification and control using real data

**DOI:** 10.1101/2023.01.08.522971

**Authors:** João A. F. Brogin, Jean Faber, Selvin Z. Reyes-Garcia, Esper A. Cavalheiro, Douglas D. Bueno

## Abstract

Epilepsy affects millions of people worldwide every year and remains an open subject for research. Current development on this field has focused on obtaining computational models to better understand its triggering mechanisms, attain realistic descriptions and study seizure suppression. Controllers have been successfully applied to mitigate epileptiform activity in dynamic models written in state-space notation, whose applicability is, however, restricted to signatures that are accurately described by them. Alternatively, autoregressive modeling (AR), a typical data-driven tool related to system identification (SI), can be directly applied to signals to generate more realistic models, and since it is inherently convertible into state-space, it can thus be used for the artificial reconstruction and attenuation of seizures as well. Considering this, the first objective of this work is to propose an SI approach using AR models to describe real epileptiform activity. The second objective is to provide a strategy for reconstructing and mitigating such activity artificially, considering non-hybrid and hybrid controllers − designed from ictal and interictal events, respectively. The results show that AR models of relatively low order represent epileptiform activities fairly well and both controllers are effective in attenuating the undesired activity while simultaneously driving the signal to an interictal stage. These findings may lead to customized models based on each signal, brain region or patient, from which it is possible to better define shape, frequency and duration of external stimuli that are necessary to attenuate seizures.

**Author summary:** Epilepsy is perhaps one of the most studied brain disorders and it is still not sufficiently well understood. The use of computational models is useful in this case since several simulations can be run using them, such that experience and insight about seizures can be gained without necessarily carrying out experiments. These models are usually designed with or without some knowledge about the brain region or phenomenon. Seizure attenuation approaches have been proposed for the first case, but they are limited to the type of seizure correctly described by the model. The present work proposes a similar procedure for the second one (where only the data are available and nothing else is assumed), which is regarded as more realistic due to its direct application on the signals and can lead to customized models for each activity, brain region or patient, defining important information such as the shape, frequency and duration of the external stimuli that must be applied to mitigate a seizure.

## Introduction

Epilepsy affects millions of people worldwide [1, 2], leading to several implications that reduce patients’ quality of life [5], which defines this disorder as a very sensitive and relevant topic of research. From the neurophysiological point of view, it is associated to the progressive hypersynchrony of neural firing over time, generated by endogenous or exogenous agents [1, 6]. However, the precise triggering mechanisms that that disrupt a healthy brain leading to a seizure are not sufficiently understood yet [7]. Currently, different papers have been focusing on the development of computational models able to identify the beginning of seizures and their main time signatures [8, 9]. These models help to explain experimental observations involving seizure onset and propagation [8, 10], mechanisms of seizure synchronization and control [11–13].

From the physical point of view, epilepsy is a multivariable and multiparameter dynamic system, described as differential equations or state-space notation [8, 13]. The parameters in these equations intend to represent biophysical factors that, if modulated, may change its overall behavior. Computationally speaking, by setting them, it is possible to pre-define the response of the system and analyze several scenarios (i.e., how the response is influenced by different parameters). In practice, if one or more of these simulations match a specific signature of real brain activity (usually given in the time domain), this implies that the set of parameters used for this case is consistent with the real activity. In system identification (SI) approaches [16, 17], this strategy is known as parameter estimation [18].

SI is a widely studied topic in some areas of knowledge, such as structure and control engineering [16, 17], where each case depends on the knowledge about the system. For example: there exist white-box models, which rely on having prior knowledge and deterministic equations that represent it [19]; gray-box models, which rely on prior knowledge and some available data (in this case, real signals from brain activity) [16, 19]; and black-box models, which only take the data into account, without any previous knowledge [16, 19].

In the case of epileptiform activity, especially involving dynamic models, SI applications in the literature are not commonly found. There exist some parameter estimation ones applied to the Epileptor model [8], whose development was an important breakthrough for a better understanding of the possible origins of seizures: Ref. [20] proposed a Baysean framework to estimate the epileptogenicity, a parameter responsible for the excitability of the model; Ref. [21], in turn, introduced a successive optimization approach to estimate all of its parameters considering uncertainties and noise. A limitation of the latter approach is that it considers only a reference output of the local field potential (LFP), not real signals. Furthermore, the application is restricted to the Epileptor model, where seizures can be accurately represented.

Works involving linear or nonlinear autoregressive models (AR/NAR), typical black-box modeling tools, have been mostly proposed in the literature for seizure detection, prediction and synchronization [22–26], but approaches considering system identification have also been used to describe the so-called NAR fingerprints of patients [27, 28]. However, the latter ones do not provide a systematic procedure to chacterize the AR equations in terms of dynamic models. If this is done, seizures can be described in state-space notation and used for purposes of control [13], for example.

Considering this, the aim of the present work is twofold: (1) to propose an SI approach, which comprises the use of real brain activity, recorded from human brain slices; (2) to provide a strategy for reconstructing real signals and applying control inputs to attenuate the epileptiform activity based on state-space plants. For the first one, AR models [29–31] are used, making the identification process simpler, since they only take into account the output of the system, which is often the case in practice.

For the second one, state-space (SS) models are obtained based on rearrangements of the AR equations [30] and converted into continuous-time models [37, 38]. Once the SS plants are available, they make it possible to design both observers and controllers^1^ for purposes of reconstructing the signal over time and then applying inputs to attenuate the undesired activity, respectively. In this sense, a seizure can be mathematically represented according to each specific signal and activity (named in this work as *customized models*), and computationally mitigated through the simulations.

Furthermore, the second objective of this work is analyzed considering two other different cases: the SS plant used to design both observer and controller is obtained from a signal undergoing epileptiform activity (named here as periodic ictal spiking − PIS) and the control inputs are applied to the same signal; the plant is obtained from a signal that represents interictal activity (named here as II) and applied to the PIS signal, which is taken as a hybrid controller in this work. The main purpose is to assess the possibility of designing observers and controllers without necessarily having access to seizure events, only the interictal ones. This is regarded as an interesting and important study in the sense that one does not need to rely on awaiting seizures to occur to be prepared for it. Although these events are cyclic [8, 39], they are often unpredictable [40, 41], leading to interictal periods that are typically much longer than the ictal ones [42], so hybrid controllers may compensate for the possible lack of PIS activity recorded.

The electrophysiological data used in this work were recorded from slices of hippocampal subfields^2^ that were surgically resected from patients with pharmacoresistent temporal-lobe epilepsy (TLE) [50]. The results show that AR models can, indeed, represent real epileptic seizures fairly well, which is particularly helpful to provide customized models for each patient or time signature. The observers can accurately recreate the real activity over time based on the identified SS plant converted from the AR model, regardless of the approach adopted (hybrid or non-hybrid). The controllers are capable of attenuating the amplitudes of the undesired spikes (representative of periodic ictal spikes) while simultaneously driving the system back to the interictal condition (II), being the non-hybrid ones more effective. At last, further implications about having these patient-specific models as well as the advantages and disadvantages of the proposed approach are also discussed in detail to evaluate the contributions achieved in this work in future practical clinical applications.

This article is organized as follows: the Methods section presents all the concepts and techniques employed for the proposed SI approach, divided into three stages: design, simulation and comparison; the next sections, Results and Discussion, comprises all the results yielded during these stages, as well as the main contributions and limitations intended in this work; at last, the Final Remarks section summarizes the main findings obtained.

## Materials and methods

This work comprises 3 major stages: 1) design, 2) simulation and 3) comparison. For the first one (1), the SI strategy is carried out to obtain the AR models, convert them from discrete state-space (DSS) to continous state-space (CSS) models and design both observers and controllers. Design is composed of 10 main steps: *i*) obtaining raw signals, which are the brain activity directly measured during clinical procedures (either II or PIS); *ii*) signal processing, which in this case consists of filtering and downsampling; *iii*) frequency analysis; *iv*) AR model definition and existence; *v*) AR model order selection and fitting; *vi*) model extension; *vii*) discrete state-space definition (DSS); *viii*) convertion to continuous state-space model (CSS); *ix*) design of the observer; *x*) design of the controller.

Simulation leads to 2 other steps: *i*) applying the observer to reconstruct the real signal, which is a copy of the PIS/II activity; and *ii*) applying the controller to attenuate the epileptiform activity while driving the system (i.e., brain region) to the II condition, where no seizure takes place. This stage is performed such that the estimated states are used as input variables for the controller. At last, the comparison stage comprises comparisons between the controlled signal (i.e., mitigated PIS driven back to II) and the original II one. For this purpose, the Euclidean norm, spectrograms/power spectral density (PSD), cross-correlations, principal component analysis (PCA) and appropriate statistical tests are considered. In this way, a comprehensive time-frequency analysis is carried out to assess the efficiency of the controller.

### Ethics Statement

The present study was approved by the Ethics Committe of Universidade Estadual Paulista “Júlio de Mesquita Filho” (CAAE 60470222.9.0000.5402), thus allowing for the use, analysis and sharing of the dataset2

### Design

#### Electrophysiological *in vitro* recordings

The signals analyzed in this work were previously recorded from the granule cell layer of the dentate gyrus (DG) and in the pyramidal cell layer of the CA1, CA2, CA3, CA4 and subiculum (SUB), by using glass electrodes placed on slices of 12 human hippocampal specimens that were surgically resected from patients with pharmacoresistent temporal lobe epilepsy. This process was approved by the Ethics Committe of Universidade Federal de São Paulo (CAAE 47551015.1.0000.5505) with written informed constent from all the patients, and then published in [50]. For further details about the procedures, please refer to the same study.

#### Signal Processing

The raw signals were properly filtered to remove any uncorrelated activity (i.e., shifting baseline, spurious harmonics), as carried out in [50]. By filtering them, the resulting activity consists mainly of only epileptic spiking or bursting behavior over time. Then, a downsampling of factor 10 is applied (from 10kHz to 1kHz), which helps significantly to reduce the computational cost and time, without compromising its overall behavior.

#### Frequency Analysis

By analyzing each signal in the frequency domain through its spectrograms, it was possible to find out which frequency components are more pronounced in the brain activity, and an appropriate time window can be selected based on them, covering a range from *f*_min_ to *f*_max_. Therefore, the minimum time window that must be used for the identification process is *T* = *t*_min_ = 1*/f*_min_, leading to *N*_min_ samples. It must be stressed that this value is only the first reference. As shall be shown in the next steps, sufficiently long time windows are applied as a conservative approach.

#### AR Model Definition

AR models are equations whose current value depends on previous observations of a time series. They try to predict present states based on its time history. The classical AR model can be expressed as [16]:

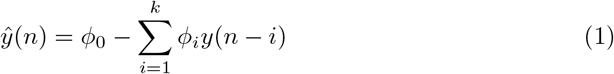

where *n* = 1, …, *N* indicates a discrete-time series, *ϕ*_*i*_, *i* = 1, …, *k*, are coefficients that weigh each past observation *y*(*n* − *i*), *i* = 1, …, *k*, of variable *ŷ*(*n*), which in this case represents the output of the system (i.e., the elecrophysiological recordings), and *ϕ*_0_ is the mean value of the time series (or the intercept).

In this work, one of the main assumptions is that the time series is stationary [30], despite local variations represented by the spikes. As shall be shown herein, with an appropriate number of AR coefficients, the epileptiform activity can be adequately reconstructed.

In practice, no AR model is capable of fully describing the dynamics of a time series, because the number of coefficients is usually limited, and for certain time intervals there might be local non-stationary behaviors. Therefore, if the error considered during the modeling is taken into account, Eq. 1 becomes:

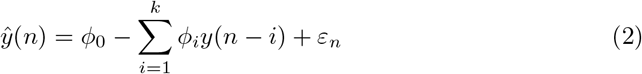

where *ε*_*n*_ is the error associated to the *n* discrete time. One possible way to evaluate model consistency is by calculating the autocorrelation function (ACF) of its residuals: [31]:

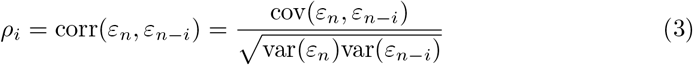

where, for simplicity of notation, *ε*_*n*_ = *ε*(*n*) and *ε*_*n*−*i*_ = *ε*(*n* − *i*), cov(.) is the covariance operator, and var(.) is the variance operator. If *ρ*_*i*_ is relatively steady and close to zero for any *i* > 0, then *ε*_*n*_ can be assumed as a Gaussian error of the type *N* ∼ (*μ, σ*^2^), where *μ* is a zero mean and *σ*^2^ is a unity variance. If this proves to be true, then the AR model has captured the significant dynamics of the time series, and the residuals are merely random fluctuations.

#### AR Model Order Selection and Fitting

Some criteria that are often used in the literature to define the order of an AR model are: the Akaike Information Criterion (AIC) and Bayesian Information Criterion (BIC). Essentially, both attempt to obtain a good trade-off between the order *k* and the sum of squared errors (SSE) between the resulting model and the original time series. Their equations can be defined, respectively, as [30, 31, 45]:

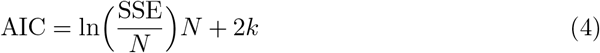

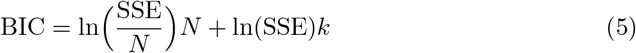

Based on these equations, when a minimum is found on the *k* vs AIC/BIC plane, it represents the best order for the model. Another possible way to assess order selection is by the mean squared error (MSE), which is closely related to the previous approach. In this work, MSE is also tested to balance the order of the model and fix a limited error, and *N* = *N*_min_ is used as the minimum time window.

#### Model Extension

This step comprises the extension of the AR(*k*) model to more samples from the measured signal. That is, if *N*_min_ samples were used to fit it, *N* > *N*_min_ samples are now included. In practice, since it is not feasible to measure an infinite time of epileptiform activity, this strategy is useful to extend the fitted model to future time instants. Naturally, this extension involves errors; however, as demonstrated in this work, they do not significantly compromise the system identification approach.

#### Discrete State-Space Definition (DSS)

Once the AR models are properly obtained, they can be converted into state-space notation according to the difference equation **x**_*k′*_ in function of the states **x**_*k*_ (which contains the dynamic matrix composed of the AR coefficients), and the measurable activity in the time domain, represented by **y**_*k*_ [30]:

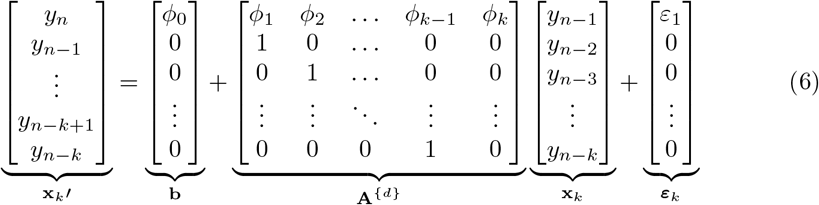

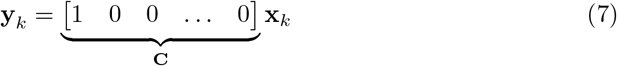

where **A**^{*d*}^ is the discrete SS model and **C** is the matrix corresponding to the measurable states in time (in this case, only the first one, *y*_*n*_, can actually be measured, whereas the remaining ones are delayed states, thus not having a specific physical meaning). Since low-frequency components are filtered out, **b** = 0. Besides, this AR modeling identifies the discrete plant and the error is evaluated a posteriori. In this sense, in practice ***ε***_*k*_ = 0 is adopted for design purposes.

#### Convertion to Continuous State-Space Model (CSS)

The previous step presents the AR model in terms of a difference equation, which is a common practice for a one-step ahead prediction, for example [29]. Although digitized signals are finite and taken as an ensemble of samples, a more realistic approach is to consider a continuous-time model that accounts for all time instants, as usually done in engineering applications [43, 44]. Besides, because the application proposed in this work requires observers and controllers, a numerical integration procedure is carried out in SS notation, which also justifies the need for a continuous-time plant. For this purpose, the formulation becomes [37, 38]:

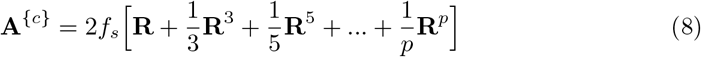

where **A**^{*c*}^ is the continuous-time model, **R** = (**A**^{*d*}^− **I**)(**A**^{*d*}^ + **I**)^−1^ and **I** is an identity matrix of the same size as **A**^{*d*}^. By adopting the Direct Truncation Method [37], the continuous-time model can be approximated, for simplicity, by:

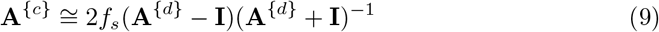

#### Design of the Observer

Suppose that, for each of the *m* = 1, 2, …, *M* time windows of size *N*_min_ presented in Fig 1, there exists an identified plant: for 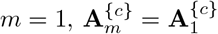 during the *N*_min_ first points; for 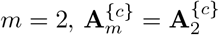 in the interval]*N*_min_, 2*N*_min_], and so forth. Then, the local models in state-space notation becomes:

**Fig 1.**
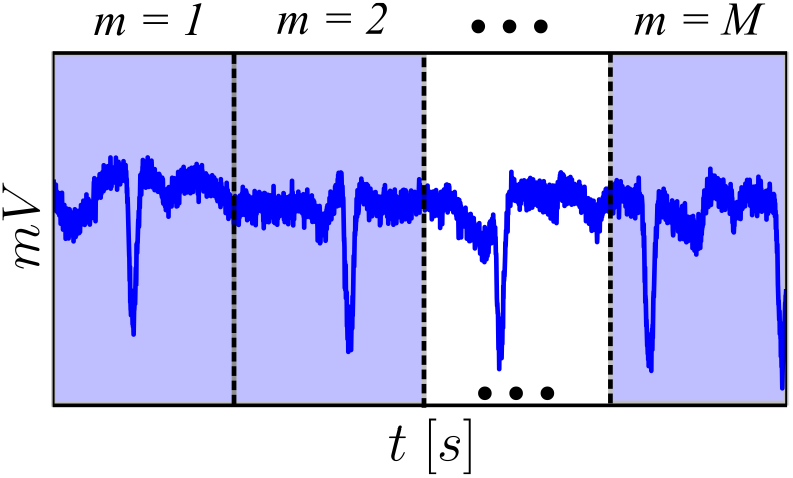
Illustration of the time windows used (whose minimum size is *N*_min_) to obtain the AR models and to which the observers and controllers are applied, ranging from. *m* = 1, 2, …, *M* . For each time window, 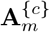 is defined and, as the observer/controller procedes in time, models are switched to adequately represent the window. The gains, however, are single for the whole signal: **G** and **L**.

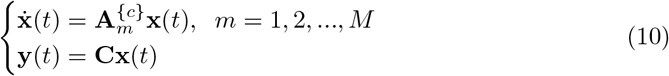

where *t* = *n/f*_*s*_ represents the continuous time variable, such that **x**(*n/f*_*s*_) ≅ **x**(*t*) [37]. This system can be understood as the mathematical representation of the real PIS/II signatures in time. However, controllers cannot be directly applied to it since the signals are only samples stored computationally, not dynamic systems. In other words, there is no access to the true system set of deterministic equations that describes the electrophysiological brain activity in hippocampal subfields or as a whole, only the signals that have been previously measured. To tackle this problem, the concept of observability needs to be applied^3^.

A state observer is commonly used when some of the states in vector **x**(*t*) do not have a precise physical meaning or cannot be measured for any reason in real applications [43]. As previously discussed, although **C** = [1 0 0 … 0]^*T*^ is consistent with the fact that only the first state is measurable and the other ones are merely artificial states, for purposes of recreating the signal in time and applying a control input afterwards, it is necessary to implement an observer such that the numerical integrator recreates its behavior artificially. For this purpose, a “copy” of the identified plant is required, represented by the following equation [43, 44]:

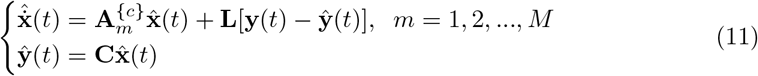

Matrix **L** weighs the relative difference between the output of the stored PIS signal **y**(*t*) and the estimated output **ŷ**(*t*) (which is in fact the recreated signal) such that, as **ŷ**(*t*) →**y**(*t*), **L**[**y**(*t*) − **ŷ**(*t*)] → 0, and the copy becomes an exact approximation of the actual output. In practice, **y**(*t*) is not a stored signal, but the real activity measured, for instance, by electrodes positioned on specific brain regions. To assess how accurate this approximation is, the error can be calculated by 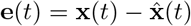 [43, 44]. By taking the derivative of this equation, the error dynamics becomes [44]:

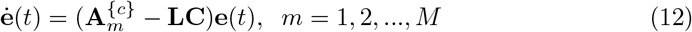

It is thus expected that **ė**(*t*) → 0 as *t* →*mN*_min_*/f*_*s*_, according to each *m* = 1, 2, .., *M* . Although the time windows switch over time, it is an interesting strategy to obtain a unique **L** that is capable of observing the whole signal, i.e., one that is valid for all time windows. To do so, it is necessary that this gain be stable across all the identified plants 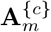. This issue is addressed in this work by taking advantage of the concept of Fuzzy Takagi-Sugeno Modeling (FTSM), which states that a nonlinear plant can be exactly reconstructed using linear submodels represented by the so-called Linear Matrix Inequalities (LMIs) [43]. In the case of the observers, they are [21, 43]:

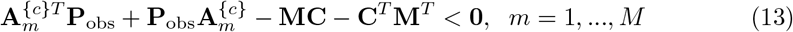

where 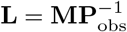 and **P**_obs_ is a positive-definite matrix [47]. If all of these inequalities are solved simultaneously, then there exists **M**, thus also assuring the existence of **L**. These inequalities are formulated based on the Lyapunov theory, so solving them implies that there exist sufficient conditions that assure stability [47]. An important difference is that, while FTSM assumes that nonlinear functions can be rewritten in terms of linear combinations between their maximum and minimum values [43], the present approach only considers the several linear models obtained from the CSS AR indentified models. It is expected that if **L** exists, it can be applied to the whole signal observing it every *m* time window as the models switch over time.

#### Design of the Controller

Considering again the *m* = 1, 2, .., *M* time windows observed and recreated by the responses of the observer, a control input can be applied to each of them according to the following equation^4^:

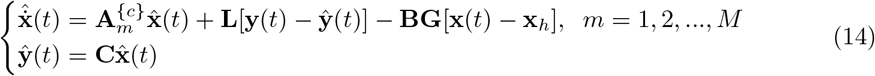

where **x**_*h*_ is the seizure-free state, that is, the condition to which the system must be driven to suppress an on-going seizure [13]. If **x**_*h*_ = 0, then the controller supposes the stability point is a zero baseline; otherwise, if **x**_*h*_ ≠ 0, it attempts to drive the current state to the desired condition while simultaneously mitigating the undesired epileptiform activity. In this work, **x**_*h*_ is taken as a reference signal in the II condition, with the first state being one of its samples at each time instant and the remaining ones equal to zero. Moreover, matrix **B** is related to the control input and can be assumed according to the design requirements or applications. To be consistent with the output matrix **C**, it is considered as **B** = diag {*B*_11_, 0, 0, …, 0}, which means that the only target state (the one that actually receives an input) is the first one, whose magnitude can be conveniently adjusted [13].

Similarly, the design of the controller can also be carried out considering a single **G** for all of the *m* = 1, 2, …, *M* time windows over time, thus assuring stability based on the Lyapunov criterion. The LMIs for the controller are [13, 43]:

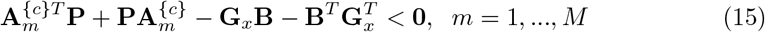

where **G**_*x*_ = **GP**^−1^. Considering that the system of interest in this case where physical inputs are intended to be applied is a brain region, close attention must be paid to the intensity of the gain **G**, so that it is properly delivered to suppress only the epileptiform activity with an acceptable input intensity [13]. This can be carried out considering a concept from FTSM, namely decay rate, defined as follows [13, 43]:

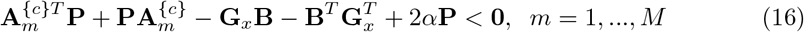

The decay rate *α* influences the positiveness of **P**, thus forcing the optimization to look for stronger/weaker gains [13, 43]. In this work, a good trade-off between this parameter and **B** is analyzed. This set of inequalities, as well as the previous ones for the observer, can be simultaneously solved using specific optimization tools, such as YALMIP [48], for MATLAB, or PICOS [49], for Python. Without loss of generality, the computational package adopted in this work is PICOS.

#### Hybrid and Non-Hybrid Observers and Controllers

This work investigates the possibility of not only designing observers and controllers based on the epileptiform/PIS activity to suppress PIS activity, and further considering the II activity applied to PIS activities. This is regarded as an interesting study in the sense that one does not need to rely on awaiting seizures to occur to take action. The two types of designs carried out are: non-hybrid (design performed using PIS to suppress PIS) and hybrid (design performed using II to suppress PIS), respectively. Essentially, this implies that the internal dynamic structure of 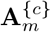 changes according to the signal from which the AR models were obtained.

### Simulation

#### Applying the Observer

Once the gain of the observer **L** has been computed and stored, it can be applied during the numerical integration process. When the signal is used for observation it represents the output **y**(*t*) of the output of the system being measured (i.e., epileptiform/PIS activities that are stored or recorded online from electrodes in practice). Therefore, the copied signal (PIS) **ŷ**(*t*) is reconstructed over time **ŷ**(*t*) → **y**(*t*) as **L**[**y**(*t*) − **ŷ**(*t*)] → 0.

#### Applying the Controller

In parallel of the **G** processing and storage, an II activity is also stored to be used as a reference **x**_*h*_. Then, as the PIS signal is observed over time, a control input **F**_*c*_ is applied to it in order to attenuate the undesired epileptiform spikes. At the same time, the controller introduces a second type of input based on **x**_*h*_. At the end of the simulation, it is expected that the reconstructed signal **ŷ** with the control input tends to the reference II signal: **ŷ** → **x**_*h*_.

### Comparison

To assess how close the reconstructed controlled signal is to the desired reference (**ŷ**→ **x**_*h*_), some criteria are adopted in this work. The first one performs a pointwise comparison between both signals according to the Euclidean norm. Then, spectrograms and PSDs are applied, which are traditional tools in engineering applications to analyze the frequency content of signals. Considering this, it is expected that the Euclidean norm ∥**ŷ** −**x**_*h*_ ∥→ 0 and the frequency contents from both signals are similar to each after the controller is activated. Furthermore, the normalized cross-correlation is applied in the time domain; this procedure aims to analyze the maximum correlation value between two signals at different lag positions. At last, PCA is carried out using the previously obtained PSDs to provide a more robust evidence that controlled signals are closer to II ones whereas uncontrolled signals are closer to PIS ones. The characterization of similarity is performed using the Kruskal-Wallis test at a 95% confidence level, and post-hoc Dunn’s test: if statistical significance can be inferred, the signals are considered to be different; if no statistical significance can be inferred, they are taken as similar.

Fig 2 presents a flowchart with all the 3 major stages proposed in this work. Note the interdependency between observer and controller: as the first one recreates the real signal computationally, a control input is applied to the recreated artificial signal to verify whether its epileptiform activity can be mitigated. This integration approach has been proposed elsewhere, but for the Epileptor model, and consistent results were obtained [21]. It must be emphasized that the present approach is only carried out because there is no access to the true system. In real applications, a control would not be applied to an artificial signal, but to a real brain region. It is thus expected that the simulations reflect real conditions sufficiently well to reproduce the results in clinical environments.

**Fig 2.**
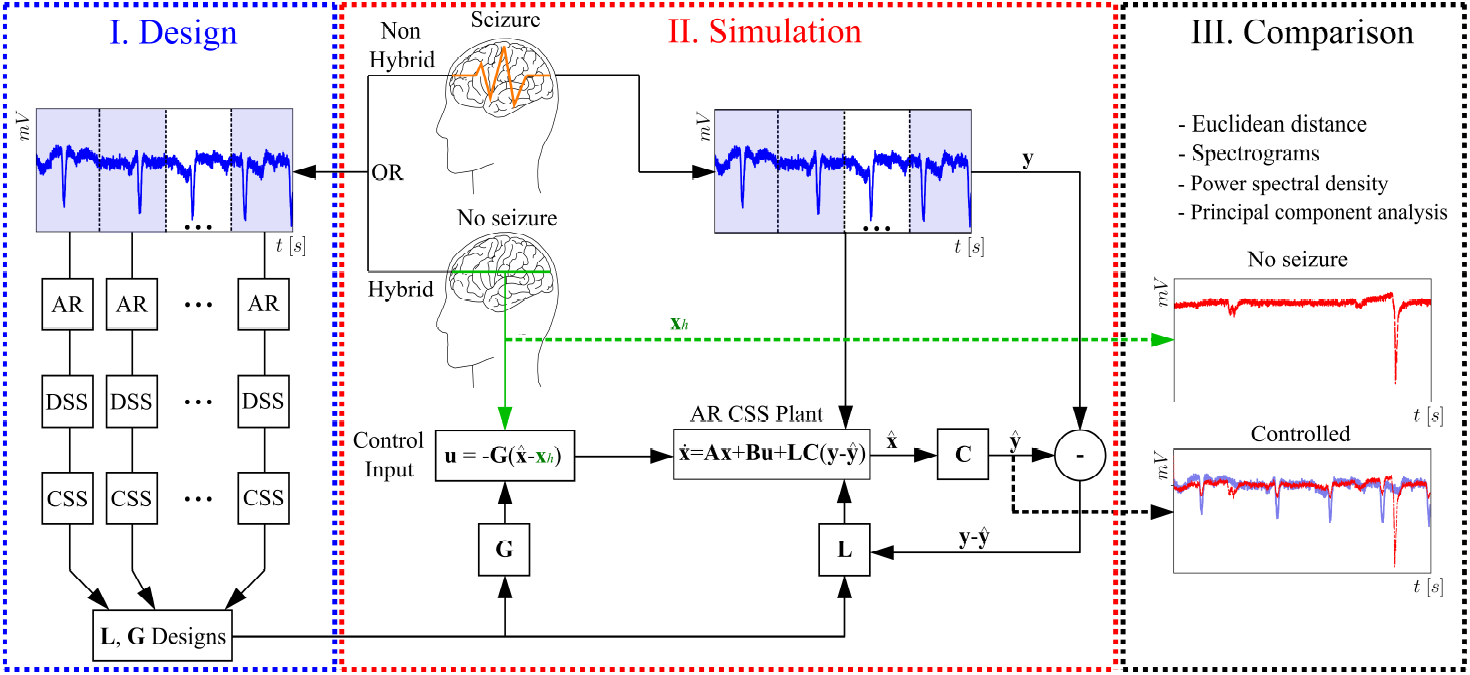
General flowchart with the 3 major stages of the proposed SI and control approach. Design: PIS or II signals are obtained beforehand (thus defining the non-hybrid or hybrid approach) so that the AR modeling can be applied; several AR models are obtained from evenly-spaced time windows, whose size is defined during steps *iii* and *iv*; the models are rearranged into DSS and then converted to CSS; the same number of plants for designing the observers and controllers is used to generate single individual gains: **G** and **L**, respectively (it is important thus to emphasize that: several models from one single lead to single gains). Simulation: II signals (**x**_*h*_) are taken as the reference behavior to which drive the system after the control is activated (i.e., no seizure condition); the real PIS activity is reconstructed based on the output matrix **C** (identifying the measurable state) and the relative difference between measured (**y**) and estimated (**ŷ**) states, weighed by the gain **L**; with the gain of the observer **G**, control inputs are applied to mitigate the PIS activity. Comparison: the controlled activity (**ŷ**) is compared to the reference behavior (**x**_*h*_) using specific techniques: Euclidean distance, spectrograms and PSDs, normalized cross-correlation and PCA. Matrices **A** and **B** correspond to the identified CSS AR (which varies over time) plant and control input one (identifying the controlled states), properly defined in the next subsections.

## Results

This section comprises the results from all stages involving the system identification approach proposed in this work, organized in subsections. For simplicity, some of them have been merged. Besides, for purposes of better explaining each of the steps, the analysis is carried out using only 2 signals, one from each condition: PIS and II. Nonetheless, other results consdering more signals can be found in Fig C in S1 text.

### Design

#### Raw signals and filtered, downsampled signals

Fig 3 presents examples of different signals obtained from the same hippocampal subfield (in this case, the subiculum− SUB) in two conditions: interictal (Fig. 3A) and ictal states (Fig. 3B). For the raw signals, there is a moving baseline over time, which is likely due to an external source of voltage that is not inherent to the spiking activity. This issue is addressed by implementing a high-pass filter, which in this case was at 0.5Hz.

**Fig 3.**
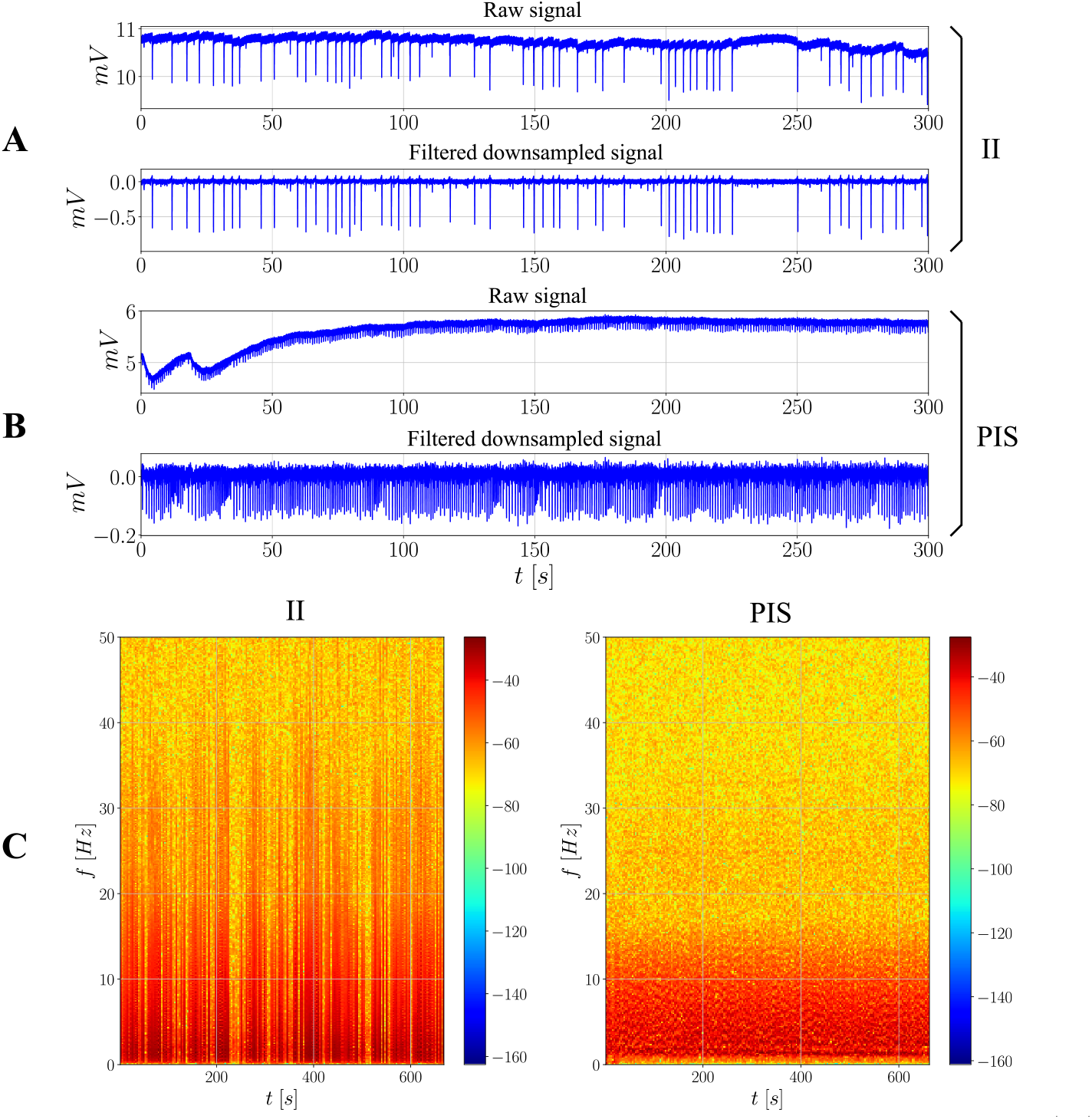
Raw and filtered downsampled signals with their respective spectrograms.(**A**) and (**B**) contain examples of raw and filtered downsampled signals from SUB in the II and PIS conditions, respectively. For the filtering, a high-pass first-order filter is applied at 0.5Hz. For the downsampling, a factor of 10 is applied in this case, which means that the initially 10000Hz-sampled series becomes a 1000Hz-sampled series. (**C**) presents the spectrograms for the same signals, both obtained after filtering and downsampling. A higher range of frequency is reglected because it does not contain further strong frequency components.

This process is not only important to unveil the spikes and bursts, but also necessary so that the mean properties of the signal are better kept over time, which is a condition for the AR models to be properly estimated [30]. Although the downsampling effect cannot be explicitely seen, a factor of 10 was applied to all signals, such that the sampling frequency becomes 1000Hz instead of the previously defined one, equal to 10000Hz.

The main difference between ictal (PIS) and interictal (II) spikings concerns the interspike intervals: while the PIS activity presensts more regular evenly-spaced spikes, the spikes of the II activity are more irregular, switching from local bursts to isolated spikes over the whole signal. A previous statistical analysis and classification of the signals used in this work has been carried out [50]. Essentially, according to the authors, the patterns exhibited in PIS signals indicate a more deterministic activity and underlying mechanism, whereas II signals suggest a more complex/chaotic one. Further considerations on this matter are out of the scope of the present work, but some of these findings are important and shall be recovered in the next sections.

#### Frequency analysis

Fig 3 also shows the spectrogram of both signals analyzed above, after filtering and downsampling (Fig. 3C). There is an increase in terms of power for the ictal activity over the whole period of time in question, which is evidenced by the darker pixels. However, in terms of frequency components, their ranges remain relatively the same for both conditions, comprising approximately from up to ∼40Hz. A higher range of frequency is not shown in the same figure because it does not contain strong components.

As a conservative approach, it is assumed in this work that: *f*_min_ = 0.5Hz. Therefore, *t*_min_ = 2 seconds, at least, to assure that all spectral activity is properly represented during the AR model definition, which leads to *N*_min_ = 2000 points considering *f*_*s*_ = 1000Hz, the sampling frequency for the downsampled series.

#### AR model definition, order selection and fitting

Since the definition of an AR model is closely linked to its fitting, which, in turn, is based on order selection, these steps are carried out together. To define the AR model, *N*_min_ comes in handy, because it has already been calculated as an adequate time resolution. As a conservative approach, this number of points is increased by a factor of 2.5 in this work, leading to *N*_min_ = 5000. By doing so, it is also possible to comprise different features over a longer period of time, thus providing a source of heterogeneity to the model.

Fig 4 presents the results for the *k* vs AIC, BIC or MSE criteria, considering the two previously studied signals, in the interictal (Fig. 4A) and ictal conditions (Fig. 4B). Note that, as the number of *k* increases, all of the measures for the criteria decrease. Additionally, the difference between previous and past values tends to become less and less abrupt, indicating that the AR(*k*) coefficients of higher order have a weaker effect in improving significantly the quality of the model. At last, both criteria provide similar results, which is evidenced by the superposition of both curves.

**Fig 4.**
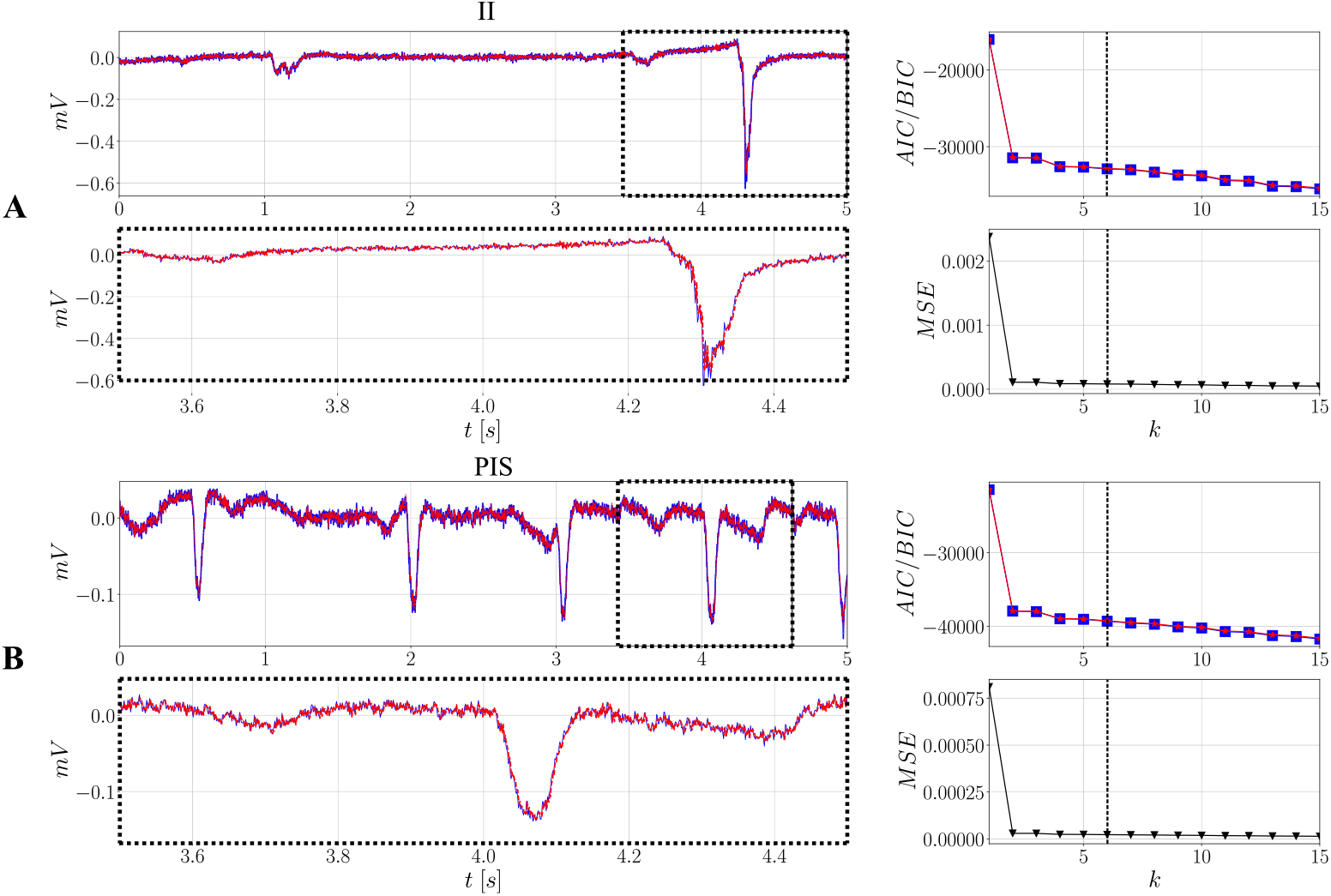
Reference II and PIS signals from SUB and their respective AR models obtained after order selection based on the AIC/BIC/MSE criteria. (**A**) contains the results for the II case whereas (**B**) constains the results for the PIS case: **–** represents the reference II/PIS signals; **- -** represents the respective fitted *AR*(*k* = 6) models. Order selection: **-**□**-** indicates AIC curves; **-**\***-** indicates BIC curves.

From the same curves, *k* = 6 has been selected for specific reasons: *i*) as mentioned, the decrease after *k* = 6 is smoother, less steep; *ii*) for higher values of *k*, some coefficients for the interictal model may assume a magnitude such that |*ϕ*| > 1, which theoretically is not adequate due to the stationarity assumption [31]; *iii*) the main goal is to keep both models as simple as possible. It is worth mentioning that, for both conditions, the overall MSE is less than 0.05%, indicating a good representation of both spiking activities.

Fig 4 also shows the models obtained for both conditions considering *k* = 6. Overall, the AR models capture well the dynamics involved in the signals, from steadier behaviors, as the fluctuations around the baselines close to zero, to the abrupt changes that arise as spikes occur. Further results concerning the autocorrelation functions of the residuals are presented in Fig A in S1 text, where similar conclusions can be drawn. Furthermore, the six coefficients found are different between the conditions. This result is important because it provides evidence that relatively simple models, as autoregressive ones, might be an interesting option to describe and distinguish types of seizures, thus providing patient-specific models with their own features.

### Model Extension

Once the AR(*k*) has been obtained considering *N*_min_, the same equations can be applied to the remaining samples of the whole time series (*N* > *N*_min_). Fig 5 presents both AR models fitted to their respective signals (interictal and ictal), as well as their MSE over time. To obtain this error, equally spaced moving windows of size *N*_min_ were taken as references so that MSE could be calculated for each of them.

**Fig 5.**
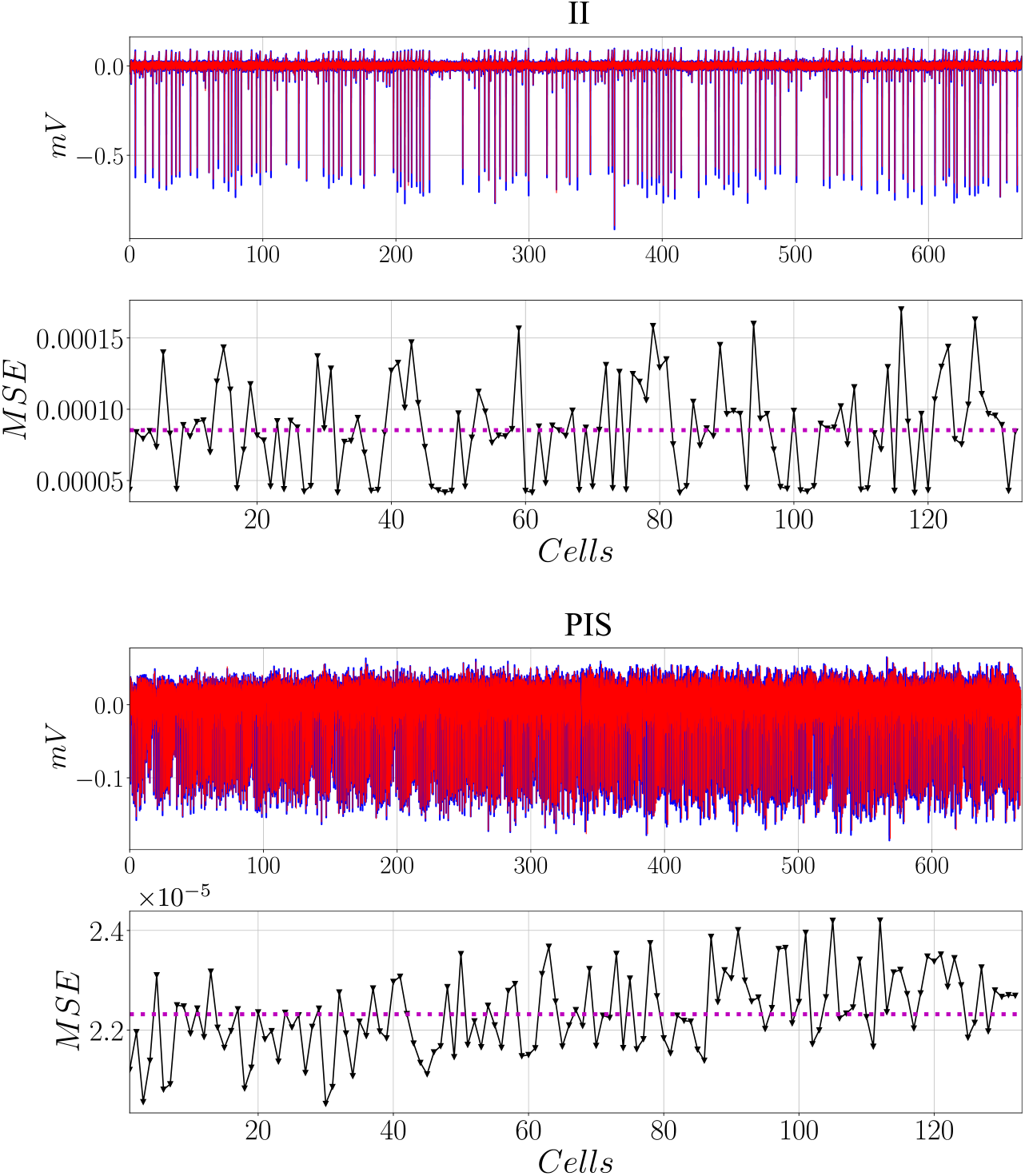
Model extensions calculated for the whole time series based on the AR models found using the first *N*_min_ samples. – indicates the filtered downsampled signals; **- -** indicates the fitted models; **…** indicates the mean values of the MSE. Each cell contains *N*_min_ samples.

The errors exhibit a quite regular behavior, whose means are below 0.01%, the value that was previously found during order selection. This result is important especially envisioning more realistic practical situations, where storing an infinite number of samples is not feasible. Therefore, epochs of interictal or ictal signatures can be acquired to be used as templates for the AR models, and then used for the upcoming brain activities.

### Discrete/Continuous State-Space and Design of the Observer/Controller

Fig 6 presents the matrices for the discrete-time models (Eq. 6), obtained directly from the AR coefficients, and the continuous-time models (Eq. 9), after the appropriate conversion. For illustration purposes, only the *m* = 1, 2, *M* models are shown, i.e., the ones obtained from the two first time windows and the last one. Light colors indicate that the magnitude of the element inside a matrix increases positively, whereas cold colors indicate this magnitude increases negatively.

**Fig 6.**
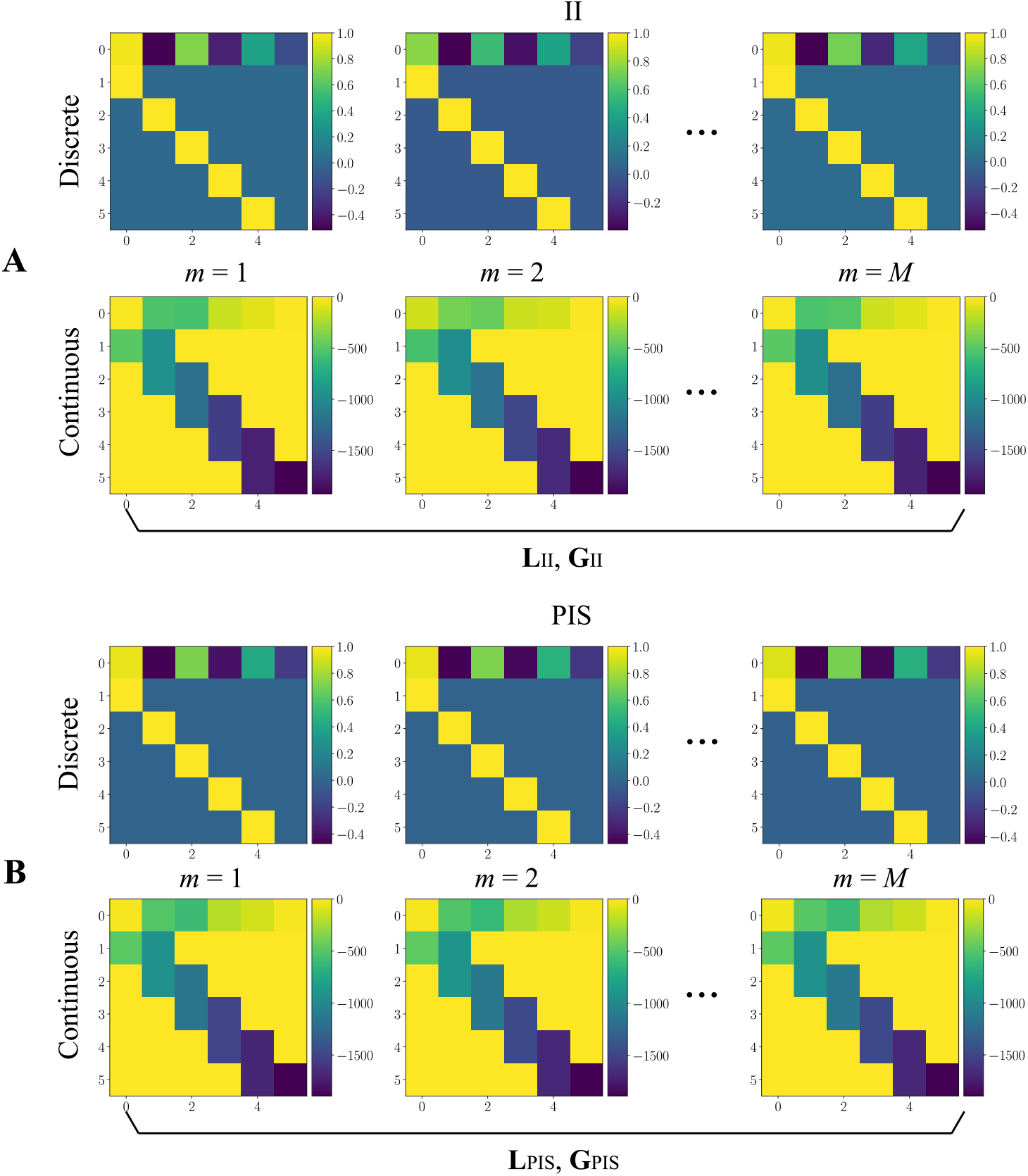
Discrete and continuous dynamic matrices obtained from the AR coefficients in state-space notation. (**A**) and (**B**) represent 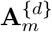 and 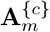, from Eqs. 6 and 9, respectively. The set of matrices 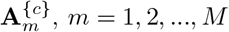, is used to compute single gains for the observer (**L**) and controller (**G**) in one of the two conditions (II or PIS). For illustration purposes, only the *m* = 1, 2, *M* models are shown, i.e., the ones obtained from the two first time windows and the last one.

All *m* = 1, 2, …, *M* together in the continuous form are used to design the gains of both observer and controller for one of the proposed schemes: hybrid (**L**_*II*_ and **G**_*II*_, Fig. 6A) and non-hybrid (**L**_*P IS*_ and **G**_*P IS*_, Fig. 6B). In this way, single gains assure dynamic stability when applied to the whole signal. Note that their structure is quite similar to each other, regardless of the condition, mostly having magnitudes of the type |*ϕ*| < 1 (for discrete models), which is desired in the case of AR modeling [31].

For clarity of understanding, Table 1 shows the AR coefficients for some of the aforementioned matrices. There are differences from the coefficients calculated in both conditions, which indicates that AR models can capture the dynamic transitions inherent to the real signals, which is reflected on their coefficients. This distinction is particularly evidenced by the fluctuation in the values between each condition, especially for II, where the first term (*ϕ*_1_) is less stable across the windows presented (characterizing the irregular pattern of II compared to a more regular PIS activity [50]). This characteristic is specific to this type of modeling and has been used in pattern recognition approaches involving, for instance, structural health monitoring [51–54] as well as analysis and validation of computational models in describing II and PIS activities [50], since the AR coefficients are sensitive to changes in the signal dynamics [52].

**Table 1.** Coefficients for AR(*k* = 6) models obtained for the SUB signal in the interictal (II) and periodic ictal spiking (PIS) conditions. Means are calculated considering all of the time windows (*m* = 1, 2, 3, …, *M*), not only the ones presented. Each line corresponds to the first line of the matrices (discrete case) from Fig. 6. The table is restricted for illustration purposes. For further results, please see Fig B in S1 text.

| Time Window (II) | $\phi_1$ | $\phi_2$ | $\phi_3$ |
| --- | --- | --- | --- |
| $m = 1$ | 0.96601012 | -0.51796275 | 0.71708485 |
| $m = 2$ | 0.74988617 | -0.39621265 | 0.56154335 |
| $m = M - 1$ | 0.66563797 | -0.43145048 | 0.51627098 |
| $m = M$ | 0.96324516 | -0.53397719 | 0.67803996 |
| Time Window (II) | $\phi_4$ | $\phi_5$ | $\phi_6$ |
| $m = 1$ | -0.37918536 | 0.35394271 | -0.15192228 |
| $m = 2$ | -0.32536739 | 0.41139278 | -0.12343279 |
| $m = M - 1$ | -0.35454543 | 0.36084365 | -0.17006476 |
| $m = M$ | -0.35530265 | 0.36815283 | -0.13206332 |
| Time Window (PIS) | $\phi_1$ | $\phi_2$ | $\phi_3$ |
| $m = 1$ | 0.94856822 | -0.47273578 | 0.70742222 |
| $m = 2$ | 0.95015544 | -0.47088456 | 0.71662639 |
| $m = M - 1$ | 0.88594754 | -0.37282567 | 0.61554666 |
| $m = M$ | 0.92252666 | -0.45359902 | 0.69951633 |
| Time Window (PIS) | $\phi_4$ | $\phi_5$ | $\phi_6$ |
| $m = 1$ | -0.40025955 | 0.42473779 | -0.21686942 |
| $m = 2$ | -0.43850637 | 0.46697371 | -0.23336034 |
| $m = M - 1$ | -0.35305278 | 0.44163889 | -0.22369959 |
| $m = M$ | -0.42437682 | 0.46004082 | -0.21459317 |

In fact, Ref. [50] used the same signals present in this work, and by applying nonlinear/nonparametric classification techniques, such as logistic regression and confidence intervals, fairly good classifications were obtained. Moreover, within each condition, the coefficients are relatively bounded over the time windows (for further results, please see Fig B in S1 text), which suggests that not only the patterns of II/PIS are sustained differently, but also that the coefficients do reflect this consistency.

### Simulation

This section comprises the results from the simulation of the action of the observer and controller proposed in this work. A fourth-order Runge-Kutta was applied as the numerical integrator, considering the time step equal to the one from the real signal (that is: *dt* = 0.001s). This procedure is implemented on the Spyder platform, using Python 3.2, which is free and open-source.

### Applying the Observer/Controller

Fig 7 presents the results obtained after the application of both observer and controller. For the sake of understanding, it was divided into three parts showing the first 5 seconds of simulation (and a zoomed in pannel during the first 0.1s): for the first one (Fig. 7A), only the observer is activated while the controller is kept off. To verify the efficiency of the observer, the copy of the system is set with different initial conditions from the ones measured in the real system. It is expected that the gain **L** weighs the relative diffence between states such that both time signatures converge. This is evidenced in the same by the fact that the error **e**(*t*) → 0 in a relative short time (first 0.1 seconds).

**Fig 7.**
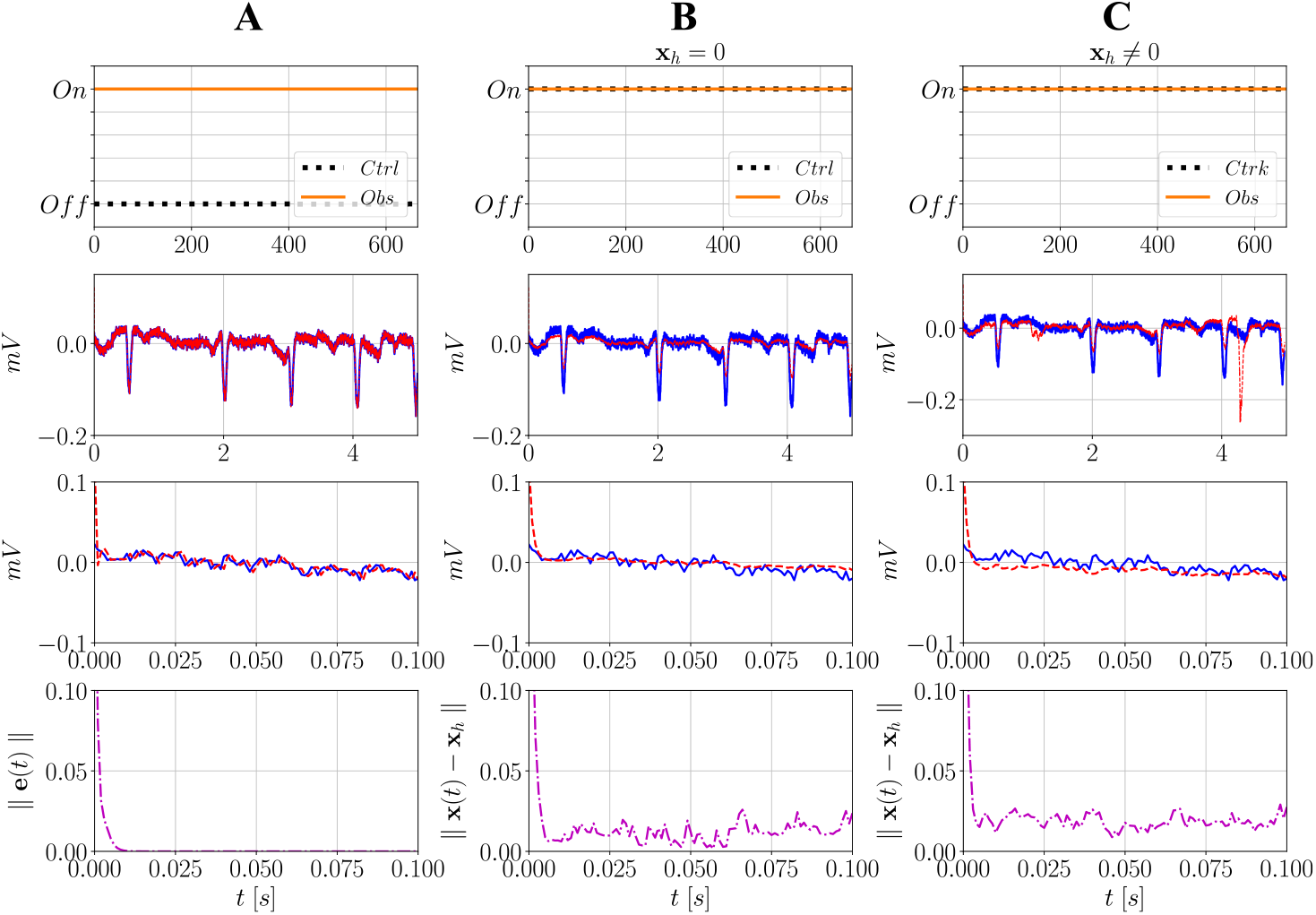
Action of the observer and controller over time and behavior of the error dynamics and Euclidean norms. In (**A**), only the observer is activated while the controller is not. In (**B**), both are activated while setting **x**_*h*_ = 0. In (**C**), both are activated while setting **x**_*h*_ ≠ 0. For the observer, **x**(0) (states of the real system) and 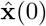 (states of the copy of the system) are set with different values and, as it progressively weighs the outputs **ŷ** and **y**, the trajectories over time converge within a limited period of time. When **x**_*h*_ ≠ 0, the controller attenuates the epileptic spikes, but attempts to mitigate them towards zero. When **x**_*h*_ 0, in turn, it delivers an additional input to simultaneously attenuate these spikes while driving the system to the II condition. In this case **L**_PIS_ and **G**_PIS_ are used

In this case, the numerical integrator reconstructs the stored signal computationally over time. In practice, however, the process of signal acquisition would be carried out from a real electrode, positioned on the brain region of interest. In other words, by designing a state observer, the real PIS/II activities can be reconstructed (or “observed”) artificially. It must be stressed that this process is different from simplyrecording the signals: if there were more states in vector 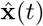, all of them would be recreated, regardless of their physical interpration. Besides, this approach enables having an artificial dynamic model where control inputs can be applied.

Fig. 7B presents the case where both observer and controller are activated from the beginning of the simulation, while setting **x**_*h*_ = 0. This implies that the control attempts only to drive the system to a constant baseline around zero. As shown, its action is efficient in attenuating the amplitudes of the spikes over time. Nonetheless, this is not the most realistic scenario, since it is desired not only to mitigate the PIS activity, but also drive the system back to the condition it was before a seizure started [13]. Thus, the reference state must necessarily be of the type **x**_*h*_ 0, as presented in Fig. 7C. Note that not only are the amplitudes of the spikes mitigated (from the PIS activity), but the controller progressively delivers the reference input to bring the signal back to the II condition.

At last, considering the Euclidean norms between the reference state **x**_*h*_ and actual states **x**(*t*), values fluctuate close to zero but do not reach this level in both applications of the controller. This is also expected, firstly because for **x**_*h*_ = 0, the difference is calculated from the signal and zero, and since a complex activity is going on, a zero baseline is never achieved. Similarly, for **x**_*h*_ ≠ 0 the spikes are attenuated but not canceled, and since it is desired that **x** → **x**_*h*_, the progressive addition of the reference **x**_*h*_ to the controller makes it interact with the PIS activity reconstructed from the observer. The final result is not an exact version of the reference, but a close one. Therefore, it is expected that the controlled signal presents a similar behavior to the reference globally, not locally. Fig 8 presents the action of both observer and controller, but now considering the whole signal from SUB. The observer remains activated all the time while the controller is only turned on at the middle of the signal (half of the entire time of simulation). In the time domain, note that there is again a clear difference in the observed signal as soon as the controller is activated. The amplitude of the undesired activity (i.e., PIS) is significantly reduced and the controller keeps delivering the input signal to drive it back to the II condition (**x**_*h*_), which is indicated by the irregular spiking pattern of higher amplitude that arises.

**Fig 8.**
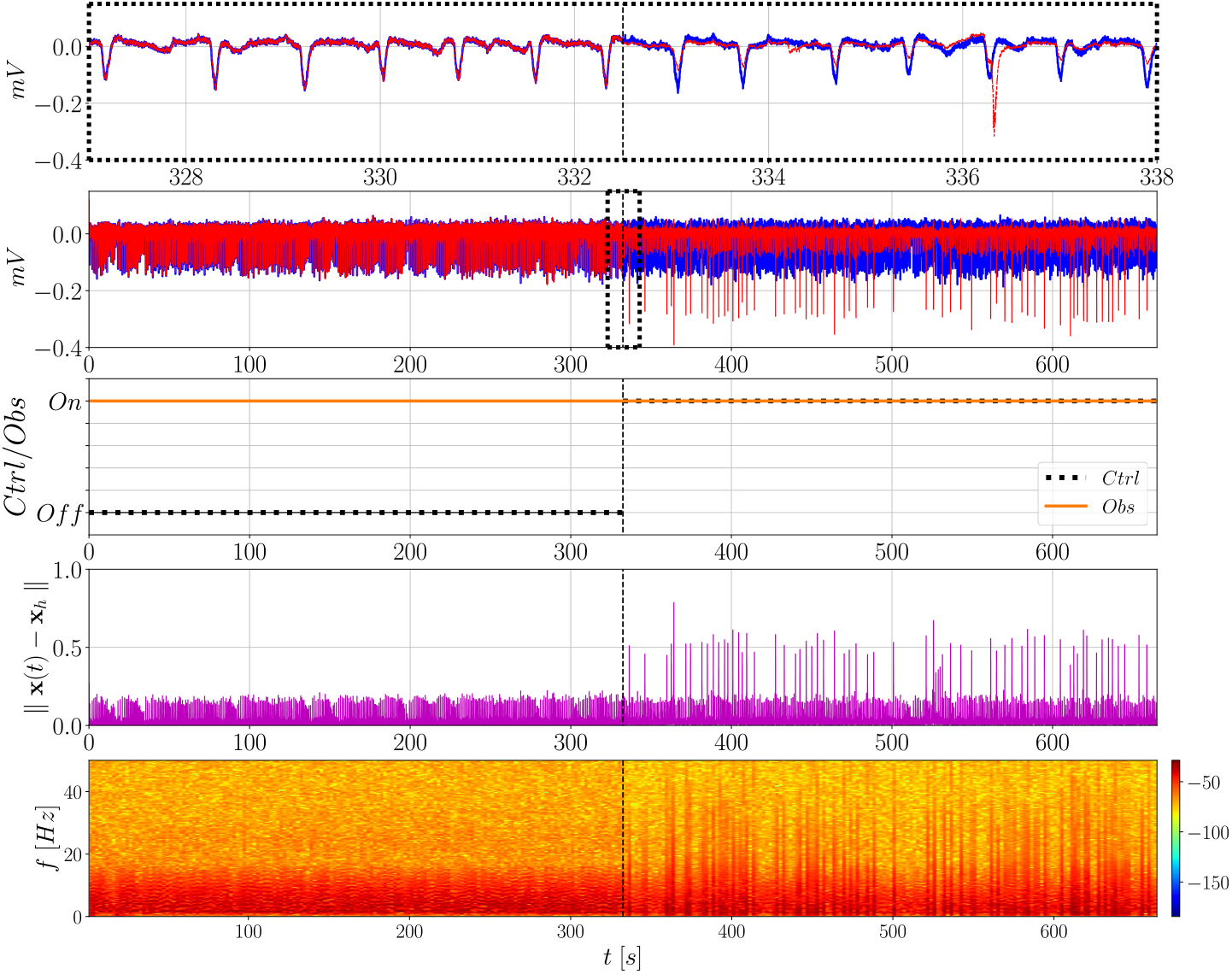
Action of the observer and controller over time analyzed in terms of the Euclidean norm ∥x(*t*) −x_*h*_∥ and frequency content over time (spectrograms). The controller is activated at half of the entire time of the signal, whereas the observer remains on during all the time. In this case **L**_PIS_ and **G**_PIS_ are used

However, such a difference is not reflected on the Euclidean norm, as previously discussed, resulting in no significant improvement. By plotting the spectrogram it becomes clear that the uncontrolled signal has a similar frequency content pattern as the PIS signal, whereas the controlled one presents higher similarity with the II signal (Fig. 3). This is the global similar behavior expected when the controller is activated.

As part of the hybrid strategy also covered in this work, Fig 9 presents similiar results, but using **L**_II_ and **G**_II_, that is, both observer and controller are designed using a SUB signal in the II condition^5^. When the controller is activated, the amplitude of the undesired activity is mitigated and the peaks of the reference signal can be seen standing out among the others, as previously. This indicates that the design can becarried out without the need of having epileptiform activity recorded, only the pre-ictal states. This is an interesting result aiming at practical applications, where seizures often occur spontaneously and unpredictably − which makes it difficult to measure them several times.

**Fig 9.**
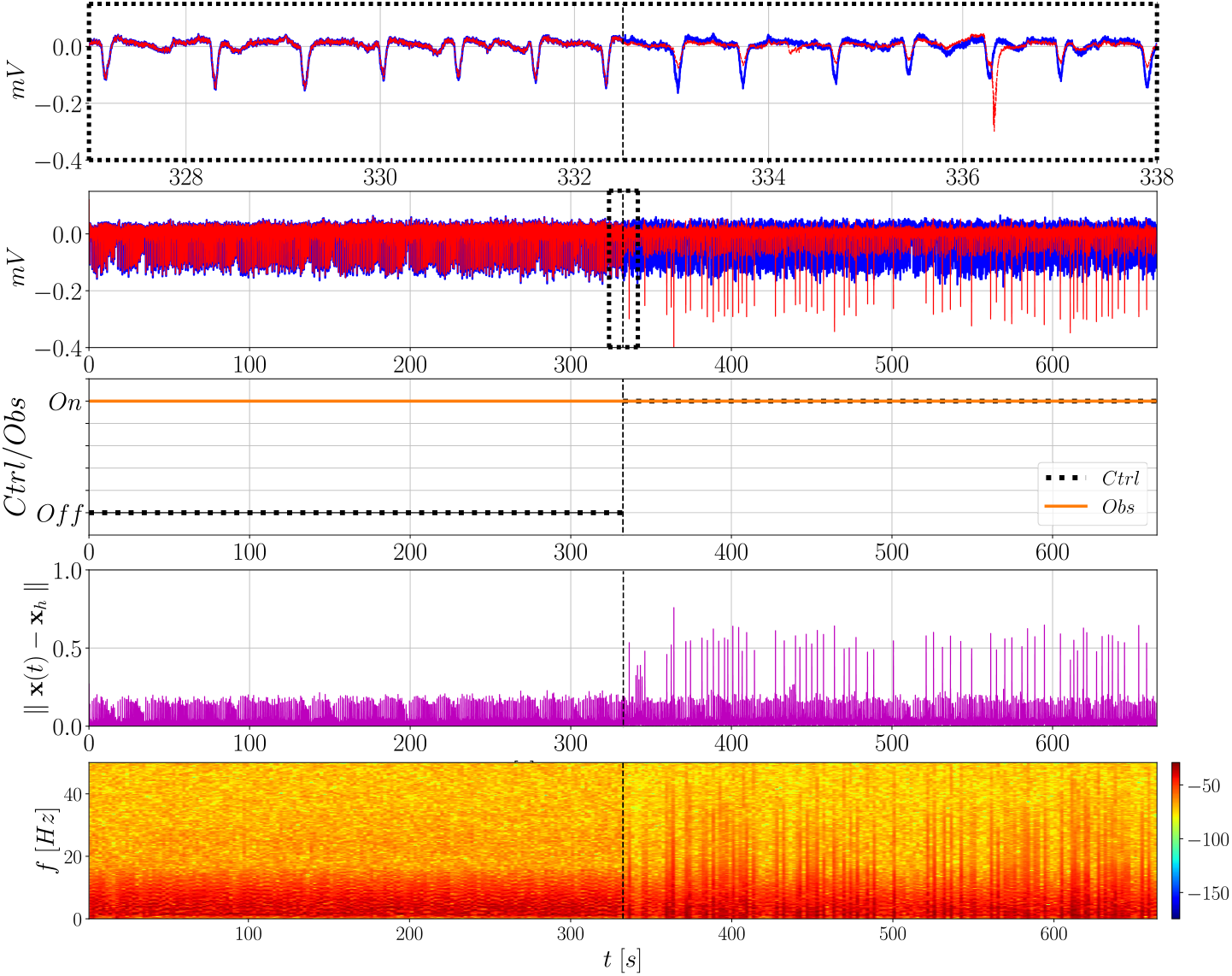
Action of the observer and controller over time analyzed in terms of the Euclidean norm ∥x(*t*) − x_*h*_∥ and frequency content over time (spectrograms). The controller is activated at half of the entire time of the signal, whereas the observer remains on during all the time. In this case **L**_II_ and **G**_II_ are used

In this work, for illustration purposes, the controller remained on for half of the duration of the signal^6^. In practice, it is not expected to act for a long time since this might not be the ideal case considering that patients would have to wear devices from which the inputs are delivered, and also for economy reasons, because if the controller is activated for short periods time, not continuously, its lifespan is increased. Besides, its intensity must also be taken into account for the same reasons. In this sense, the amplitude of such inputs must be adequately tuned so as not to compromise the health of desired brain activities. Fig 10 presents the responses of the signals in time when the parameters of the controller (*α* and *B*_11_) are tuned to decrease the intensity of the input (for both hybrid and non-hybrid cases). For illustration purposes, only the first 20 seconds of activation (and zoomed in windows) are shown. For the non-hybrid controller (Fig. 10A), when the amplitude is limited it changes from its previous highest peak (based on the time window analyzed) at ∼503.06mV to 341.43mV (in modulus), a reduction of more than 32%. Naturally, this implies that amplitudes of the undesired activity are not as efficiently mitigated as before, but a considerable attenuation can be seen, specially in the zoomed in epochs of the signal.

**Fig 10.**
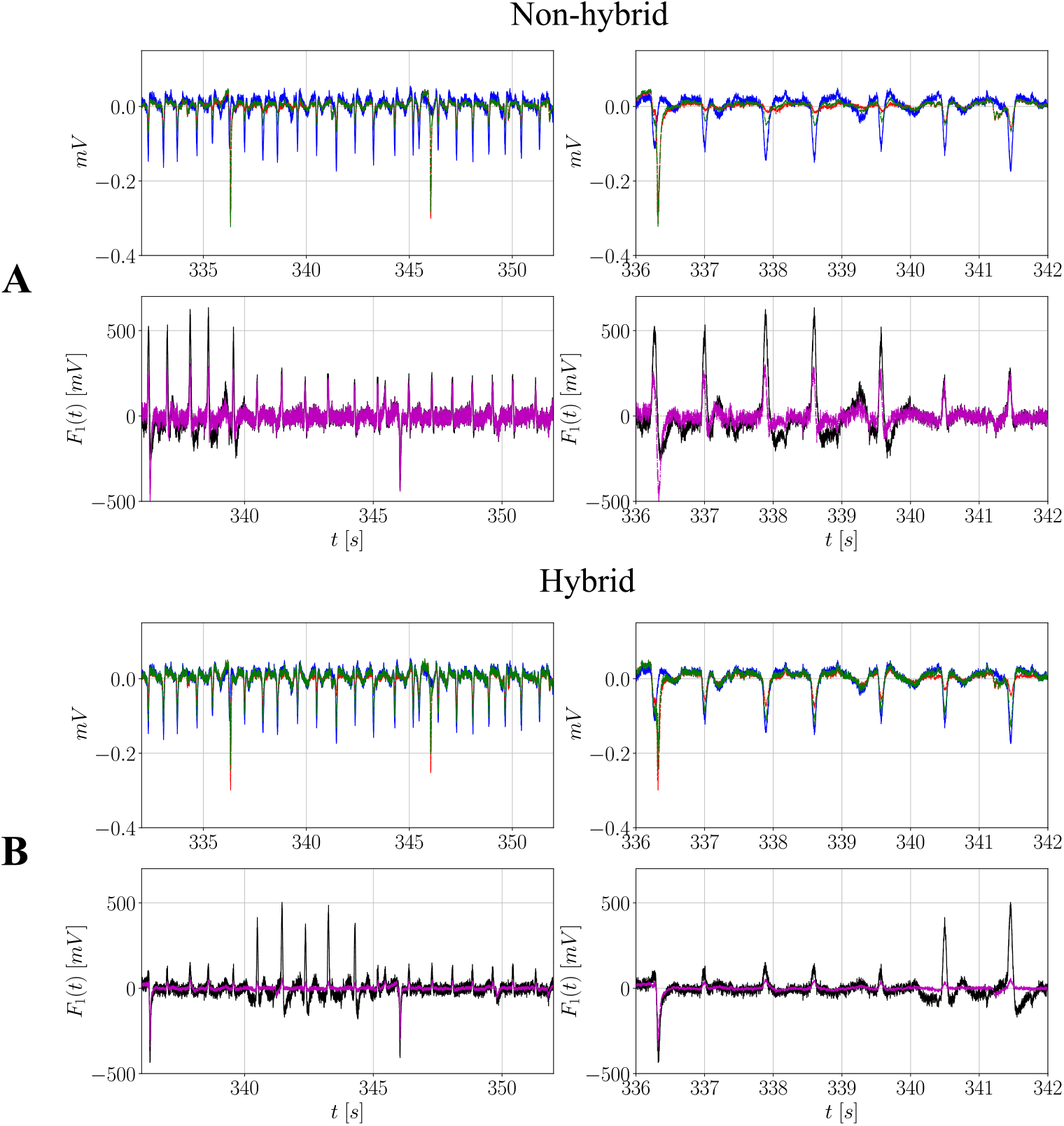
Effect of tuning *α* and *B*_11_ on the input response of the signal in the time domain. (**A**) contains results for the non-hybrid case, whereas (**B**) contains the results for the hybrid case: **–** represents the reference PIS signal (from SUB); **- -** represents the controlled signal before tuning the parameters; **-.-** represents the controlled signal after tuning them. The input forces are indicated as: **–** represents the control input without tuning and **-.-** represents the same input after tuning. In this case **L**_PIS_ and **G**_PIS_ and **L**_II_ and **G**_II_ are used, respectively. The parameters before tuning are: *α* = 40 and *B*_11_ = 8.5.10^−5^ (non-hybrid), and *α* = 20 and *B*_11_ = 7.10^−5^ (hybrid). The parameter after tuning are: *α* = 20 and *B*_11_ = 8.5.10^−5^ (non-hybrid), and *α* = 0 and *B*_11_ = 5.10^−5^ (hybrid)

Concerning the shape of the control stimulus, two components can be distinguished: the first one acts in the opposite sense of the PIS activity, presenting spikes of inverse magnitude; the second one, in turn, is responsible for delivering spikes of irregular time intervals (similar to the II activity). This finding is consistent with previous observations made in the literature, where continuous periodic stimulation/trains of pulses [55–57] or randomly-spaced ones [58] have been relatively efficient in suppressing seizures. Thus, an interesting investigation in practice would be the use of combined types of stimulation to verify their controlling effects in real epileptiform activity. As for the inputs of the hybrid controller (Fig. 10B), the main difference compared to the first one is that its amplitude decreases from ∼ 634.45mV to 504.34mV (a reduction of approximately 21% in modulus). For both of them, a significant attenuation is observed nonetheless.

### Comparison

To better assess the efficiency of the controller and endorse the previously presented results of Euclidean norm and spectrograms, this section presents results for the comparisons using PSDs, cross-correlations and PCA. For the sake of understanding: reference II represents the stored II signal (**x**_h_); reference PIS represents the stored PIS signal to be observed/controlled; the uncontrolled signal represents the observed PIS signal after activating the observer; the controlled one, in turn, represents the signal after activating the controller.

### Power spectral density (PSD) and principal component analysis (PCA)

Figs 11 and 12 summarize all of these metrics for the non-hybrid and hybrid controllers, respectively. To generate the PSDs, several *N*_min_-long windows were considered such that they are taken as random variables, which makes it possible to calculate mean PSDs with standard deviations and confidence intervals (Fig. 11A and Fig. 12A). By analyzing the means (Fig. 11B and Fig. 12B), two interesting patterns can be observed concerning the controlled signal compared to the uncontrolled one: there is a slight decrease in power up to ∼10Hz and an increase after this frequency, consistent with the fact that II represents more complex activities, thus covering a large frequency band, while PIS is a more deterministic periodic activity [50]. This suggests that upon activation of the controller, the signal tends to move towards the reference II and leave the PIS condition, regardless of the control strategy.

**Fig 11.**
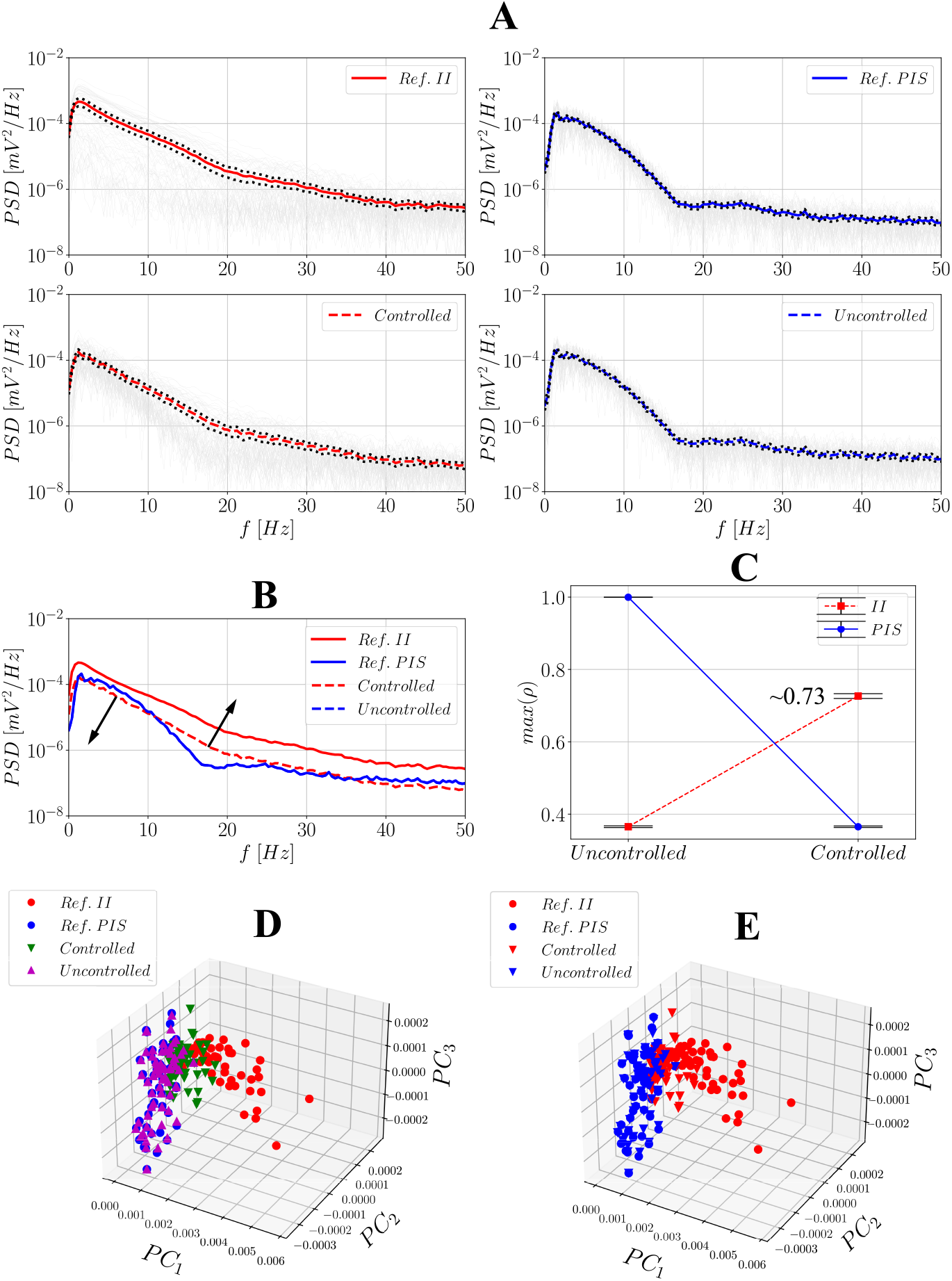
Comparison between the reference observed II and PIS signals and the uncontrolled/controlled ones using PSDs and PCA after activating the non-hybrid controller. (**A**) contains the PSD results: **–** represent the PSDs obtained for each of the *N*_min_-long windows; **–** represents the mean PSD of the reference/observed II signal; **- -** represents the mean PSD of the controlled signal; **–** represents the mean PSD of the PIS reference/observed signal; **- -** represents the mean PSD of the controlled signal; **…** represents the confidence intervals at a 95% level. (**B**) represents all the mean PSDs together for visual inspection; (**C**) contains the maximum cross-correlation values between the combinations reference II/PIS and uncontrolled/controlled signals using all of the *N*_min_-long windows and their respective confidence intervals at a 95% level. (**D**) and (**E**) present the principal components of PCA highlighting the clusters individually and after considering II/controlled and PIS/uncontrolled groups.

**Fig 12.**
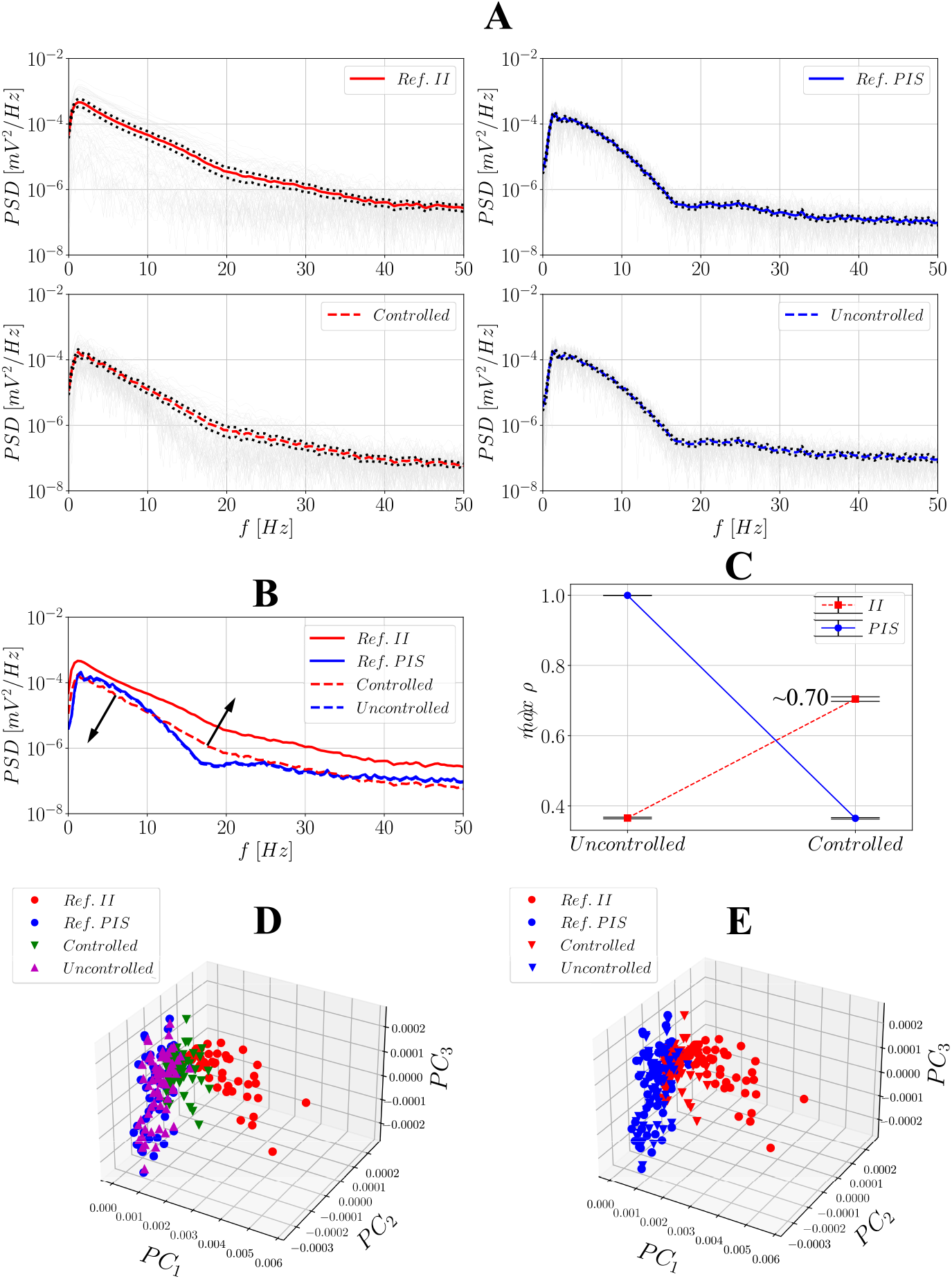
Comparison between the reference observed II and PIS signals and the uncontrolled/controlled ones using PSDs and PCA after activating the hybrid controller. (**A**) contains the PSD results: **–** represent the PSDs obtained for each of the *N*_min_-long windows; **–** represents the mean PSD of the reference/observed II signal; **- -** represents the mean PSD of the controlled signal; **–** represents the mean PSD of the PIS reference/observed signal; **- -** represents the mean PSD of the controlled signal; **…** represents the confidence intervals at a 95% level. (**B**) represents all the mean PSDs together for visual inspection; (**C**) contains the maximum cross-correlation values between the combinations reference II/PIS and uncontrolled/controlled signals using all of the *N*_min_-long windows and their respective confidence intervals at a 95% level. (**D**) and (**E**) present the principal components of PCA highlighting the clusters individually and after considering II/controlled and PIS/uncontrolled groups.

Fig. 11C and Fig. 12C show the results for the analysis of maximum cross-correlation coefficients. Each of the *N*_min_-long window was taken as a sample; then, uncontrolled and controlled signals were both compared to references II and PIS. As expected, PIS activities correlate better with uncontrolled ones, whereas II activities correlate better with controlled ones, which again evidences the effectiveness of the controller, but analyzed in the time domain − for this metric, the non-hybrid controller has a slightly better performance, with ∼0.73 vs ∼0.70. In the 3-dimensional principal component space, when each of the *N*_min_-long PSD windows is projected four different clusters are revealed, corresponding to references II and PIS and uncontrolled/controlled signals. When plotted together, regardless of the control strategy used, the samples from uncontrolled signals are spatially closer to PIS references, reference II is farther from the groups, and samples from controlled signals fall in between. When plotting only two categories (relating uncontrolled to PIS samples and controlled to II ones), the difference between these clusters becomes more pronounced.

To assess statistical similarity between these four clusters, a bilateral Kruskal-Wallis test was carried out considering only the second principal component (*PC*_2_), which proved to be the best one to distinguish them. This test yielded *H* = 156.01 and *p* ≪ 0.001 for the non-hybrid controller and *H* = 153.46 and *p* ≪ 0.001 for the hybrid one, thus indicating the samples are statistically different. Next, the post-hoc Dunn’s test was also performed to evaluate individual differences, whose results are summarized in Table 2. Again, regardless of the controller, this test can efficiently distinguish reference PIS from reference II and uncontrolled from controlled signals. Besides, it does not distinguish between reference PIS and the uncontrolled signal, and between reference II and the controlled signal at 99%, which endorses the effect of the controller in restoring the signal to the II condition.

**Table 2.** P-values obtained from the post-hoc Dunn’s test considering the samples projected on the second principal component *PC*_2_, for both non-hybrid and hybrid controllers. The following notation is adopted: *p* ≪0.001 if *p* < 10^−5^, and *p* ≪ 0.001 if *p* < 10^−6^.

| Non-hybrid | Reference II | Reference PIS | Controlled | Uncontrolled |
| --- | --- | --- | --- | --- |
| Reference II | 1.0 | $\lll 0.001$ | 0.022 | $\lll 0.001$ |
| Reference PIS | $\lll 0.001$ | 1.0 | $\lll 0.001$ | 0.940 |
| Controlled | 0.022 | $\lll 0.001$ | 1.0 | $\lll 0.001$ |
| Uncontrolled | $\lll 0.001$ | 0.940 | $\lll 0.001$ | 1.0 |
| Hybrid | Reference II | Reference PIS | Controlled | Uncontrolled |
| Reference II | 1.0 | $\lll 0.001$ | 0.012 | $\lll 0.001$ |
| Reference PIS | $\lll 0.001$ | 1.0 | $\ll 0.001$ | 0.573 |
| Controlled | 0.012 | $\ll 0.001$ | 1.0 | $\ll 0.001$ |
| Uncontrolled | $\lll 0.001$ | 0.573 | $\ll 0.001$ | 1.0 |

## Discussion

This work addressed a relevant challenge in the context of epileptic seizures, which is obtaining a data-based model that describes its signatures over time. System identification approaches involving epileptiform activity (or from interictal to ictal states) are not commonly found in the literature. An important implication of doing so is to provide models that are patient-specific or seizure-specific; that is, models that adept to each type of time signature or patient. From these models, observers and controllers can be designed for purposes of recreating the epileptiform dynamics and controlling it artificially.

From the time-frequency analysis, the total number of samples was reduced, which considerably relieves the computational burden without compromising the signals’ inherent dynamics. In addition, the main frequency components were identified and used as a reference from which to define the templates (several time windows over time) for the AR models with an appropriate fixed time length. This strategy enabled obtaining AR models of reduced order: in this work, *AR*(*k* = 6) provided a good trade-off between accuracy and complexity, resulting in low values oF MSE and other relevant criteria, such as AIC and BIC. Naturally, as the order gets higher, models become increasingly closer to their reference signals. However, one of the main goals of the proposed approach is to make the models as simple as possible (specially envisioning the later stages of state-space representation), besides avoiding potential problems of overfitting. In this sense, the models can sufficiently approximate the inherent dynamics of periodic ictal activity (PIS) and interictal activity (II), distinguishing them according to the AR coefficients obtained, thus endorsing previous results using the same signals [50].

For the model extension step, the AR(6) model calculated from the first *N*_min_ samples was reapplied to the whole signal (*N* > *N*_min_), in both conditions (PIS and II). Since the evolution of the MSE over time was stable, this suggests that the identification from restricted time intervals can be used in other parts of the signal too, a fact that is also endorsed by the consistency of AR coefficients over time windows presented in Table 1 and Fig B in S1 text. This result is particularly interesting because, in practice, obtaining arbitrarily long time signals of epileptiform activity, especially from humans, usually involves complex procedures, or recordings may not be readily available. Therefore, even with reduced information about the time signatures, it is possible to infer about their remaining dynamics over time, which is also important for the next steps of designing observers and controllers.

When applying the state observer, it was verified that the dynamic error **e**(*t*) →**0** during the first instants of the simulation. This makes it possible to recreate the time signature over time accurately from stored signals, as the case of this work. This result is consistent with a previous study that reconstructed the unmeasurable activity of the Epileptor model based on its measurable activity [21]. The main difference is that this time the reference signal is not dynamically simulated through the numerical integrator, but approximated based on observations of stored signals. In practice, it is expected that the observed signal is not a stored one, but brain activity measured in real time. As long as such activity can be properly recorded, the proposed approach can be applied.

As for the controllers, its activation could attenuate significantly the amplitude of the undesired activity, i.e., periodic ictal spiking (PIS), while simultaneously driving the system back to the interictal condition (II). To do so, it is required to record II events previously so as to provide a dataset from which to choose **x**_*h*_. This is an interesting result because, since seizures present a wide variety, the reference for each case and design is seizure/patient-specific as well. The design of hybrid controllers (state-space plant built using II signals applied to PIS signals) proved to be relatively effective, which means that it is still possible to design such a hybrid control in the absence of ictal events, a good advantage considering that seizures cannot be easily measured and often occur abruptly.

The metrics used to evaluate how close controlled signals are to their reference desired ones provided complementary results. The Euclidean norm could be reasonably applied during the first seconds of simulation to visualize the efficiency of the controller along with the dynamic error; however, for longer time windows, the norm did not prove to be the best measure for this purpose. In other words, the norm is an interesting tool to assess small-scale events, not global ones. On the other hand, both spectrograms and PSDs provided relevant information regarding the frequency content of signals: they revealed that PIS activities have higher power in lower frequency bands (∼10Hz) whereas II activities spread over the spectrum. Upon the activation of the controller, the controlled signal tends to recover a significant part of power lost in higher bands, which is also reflected on the similarity between both II and controlled signal’s spectrograms.

Concerning the normalized cross-correlation, both controllers provided significant results in the sense that PIS activities correlate better with uncontrolled signals and II activities correlate better with controlled ones. This effect is, however, slightly stronger for the non-hybrid controller. At last, PCA confirmed that both controllers yield easily separable clusters in the principal component space, and they can be distinguished statistically.

Control inputs could be tuned and constrained, which is essential in practical applications not to deliver unnecessarily strong stimuli to the brain region of interest. The shape of the input stimulus is also an important result in this work. As seen, previous approaches considering trains of pulses or continuous stimulation for purposes of seizure suppression are commonly found in the literature [55–57]. Alternative approaches considering irregular time intervals for the simulation can also be found [58]. This raises an important question on which type of simulation is the most effective. On the one hand, applying pulses of inverse magnitude to “cancel out” the spikes of PIS activities seems an acceptable premise, which may explain the similarity of the results obtained here with the first studies. On the other hand, the reasoning behind aperiodic stimulation lies on attempting to disengage the hypersynchronism and reverberating effects of neural networks [58], which is also promising considering epilepsy as a progressive hypersynchronization process [1] and consistent with the definition of **x**_*h*_.

From a physical point of view, epilepsy can be interpreted as the progressive loss of complexity due to a hypersynchronized activity [1, 50], so considering this context and the results obtained here for the control inputs, it may be inferred that periodic stimulation acts on reducing the on-going undesired PIS activity, whereas aperiodic stimulation (**x**(*t*) →**x**_*h*_) acts on progressively re-establishing the complexity of the brain region that had been oversynchronized. In this sense, combined types of stimuli remains an open topic for future investigation.

## Final Remarks

This work proposed an efficient approach for attenuating the epileptiform activity of electrophysiological recordings considering the artificial reconstruction of the stored signals based on AR modeling and an integration between state observers and controllers. Several stages/steps of design, application and comparison are involved, thus characterizing a systematic and straightforward procedure that has not been reported elsewhere.

The results provided evidence that attenuating the undesired effects of real seizures can be computationally emulated, being efficient for different hippocampal subfields (DG, CA1, CA2, CA3, CA4 and SUB). Both non-hybrid and hybrid approaches can be used for this purpose: the first one provides stronger input stimuli to mitigate periodic ictal spikings while simultaneaously driving the signal to the interictal-like behavior; the second one, despite causing slightly milder attenuation effects, is also able to generate effective outcomes. This endorses the possibility of designing controllers in the absence of sufficient PIS recordings, which is considered an advantage in this work given the inherent difficulty in acquiring signals with seizure events.

The shape of the control input seems consistent with previous approaches that adopt regular and irregular stimulation individually for seizure suppression. A major contribution in this sense found in this work is that they may be more effective if combined, applied as a superposed one consisting of periodic pulses of inverse magnitude (similar to the regular stimuli) and chaotically-behaved ones (closer to the irregular stimuli). These findings may be of great interest for real applications comprising the modeling and description of seizures (characterized by the AR models), and seizure mitigation and suppression.

## Supporting information

**S1 Appendix**. Complementary results including the autocorrelation function of the errors, AR coefficients and attenuation of epileptiform activity in DG, CA1 and CA2.

**S1 Fig. A Autocorrelation funtion** *ρ* **considering the lags of the residuals obtained after estimation of the both AR models (for II and PIS, respectively) for the first time window:** *m* = 1. The signal used in this case was from SUB, as in the main body of the manuscript.

**S1 Fig. B AR coefficients of the identified models for the II and PIS signals from SUB to design matrices** 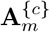, *m* = 1, 2, .., *M* **and the gains L**_**II**_ **and G**_**II**_, **or L**_**PIS**_ **and G**_**PIS**_. **–** represents the coefficients; **- -** represents their upper and lower limits.

**S1 Fig. C Performance of the controller when applied to signals coming from different hippocampal subfields: DG, CA2 and CA1. –** represents the uncontrolled activity; **- -** represents the controlled activity. In this case **L**_PIS_ and **G**_PIS_ are used without any constraints on the input. The observers and controllers were designed according to the available dataset, so results for CA3 and CA4 could not be generated due to the lack of PIS activity available for them.

**S1 Fig. D Signals used to design both gains L**_**II**_ **and G**_**II**_ **in the hybrid approach and the healthy reference x**_**h**_. In this case, signal number 21 from SUB is used for the design, whereas signal number 18 is used as a reference to which drive the system.

## Acknowledgements

To prof. Carla A. Scorza, Prof. Ricardo S. Centeno and Prof. Elza M. T. Yacubian, who contributed to the execution of previous phases of the project and the possibility of obtaining the electrophysiological recordings analyzed.

## Author contributions

Author contributions were as follows: J.A.F.B., J.F. and D.D.B. designed the research, S.Z.R.G. performed the electrophysiological experiments, J.A.F.B. implemented the computational models and with J.F. and D.D.B. analysed their data, J.A.F.B., J.F., S.Z.R.G., E.A.C. and D.D.B. wrote the paper. All authors reviewed the manuscript.

## Data and code availability

The data used in the present work will be available upon request. The codes developed for all analyzes are available on GitHub at: https://github.com/jafbrogin/identification-control-epileptiform-activity.

## Footnotes

1 These tools are commonly found in control engineering. Observers are useful for estimating abstract/unmeasurable variables from mathematical models based on the measurable ones [43, 44]; controllers, in turn, play the role of external inputs whose objective is to drive the system to a desired condition or state, thus mitigating undesired oscillations, for example [43, 44].

2 Dentate Gyrus (DG), Cornu Ammonis 1, 2, 3 and 4 (CA1, CA2, CA3, CA4) and Subiculum (SUB).

3 A similar approach has already been carried for the Epileptor model [46], where the system’s equations are known [8]. Essentially, based only on the measurable local field potential, the unmeasurable variables from the model were reconstructed [46].

4 A similar approach has already been carried for the Epileptor model too [13].

5 A different signal from **x**_h_ is used to match the amplitude of the epileptiform activity without significantly increasing the gains of the controller. Futher details can be found in Fig D in S1 text.

6 In approaches where the system’s equations are deterministic, such as the one applied to Epileptor [13], the control can be activated for a few seconds to suppress a seizure. In this work, only the signals are known, that is, the partial representation in time of a broader system, so control inputs that last longer must be applied to better visualize their effects. This is also discussed in detail in the next sections.

