## Supplemental material for "Epileptic seizure suppression: a computational approach for identification and control using real data"

Fig A depicts the autocorrelation function of the residuals of both AR models analyzed in the main body of the manuscript for the SUB signal (II and PIS, Figs. AA and AB, respectively). For both conditions, the same pattern can be observed: at the first lags, or time delays, the correlation values remain slightly higher than the confidence levels (at 95%); this indicates that these values of $\rho$ are significant. As for the correlation for the next lags, they tend to decrease in magnitude and shrink into the confidence levels, which in turn indicates that they are not significant. This behavior is expected due to the relative simplicity of the model, which cannot fully capture all the dynamics involved in the real epileptiform activity. Nevertheless, since the values of $\rho$ are low in general, the AR($k = 6$) models are taken as sufficiently good representations of the original signals.

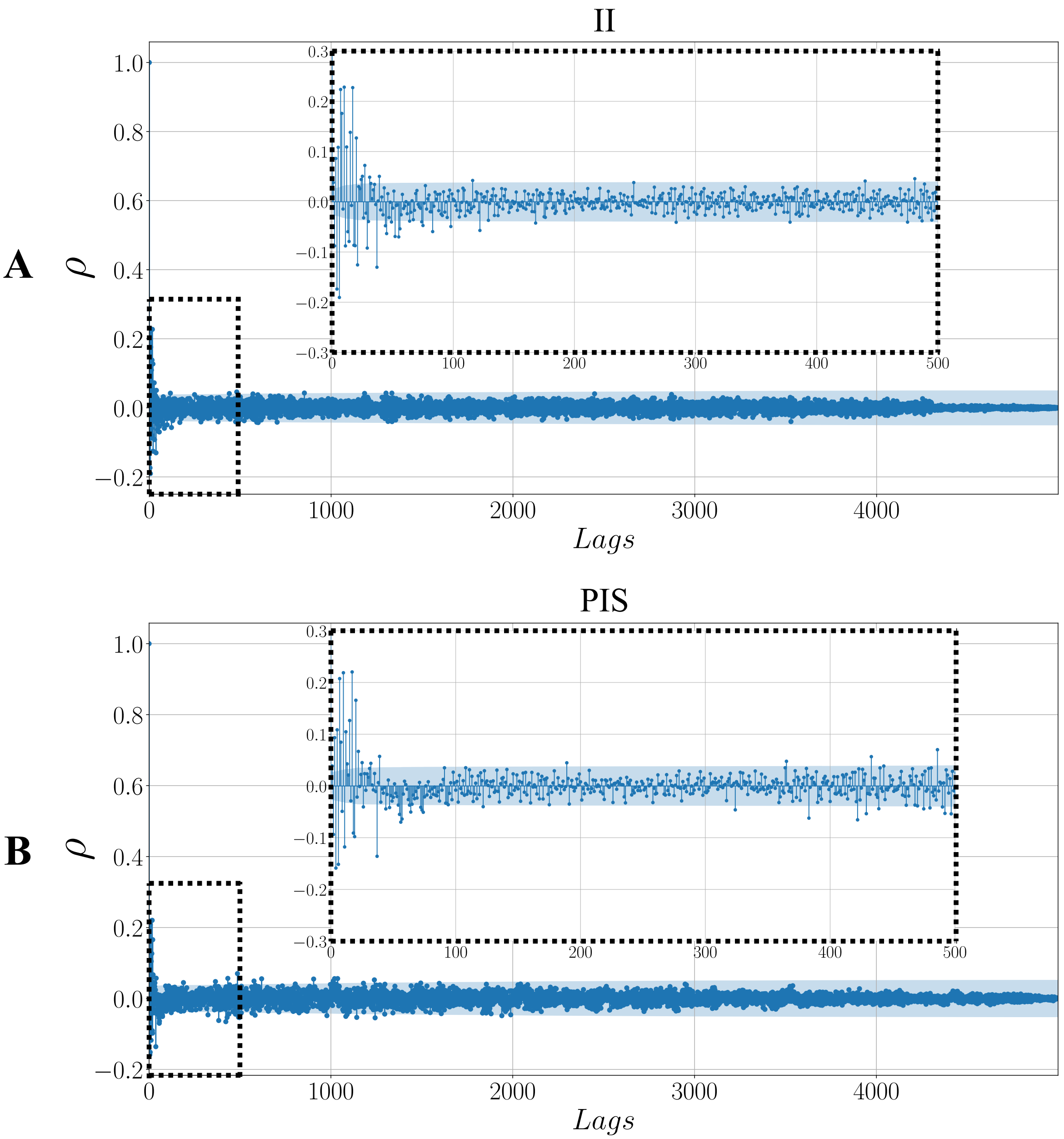


**Fig A. Autocorrelation funtion $\rho$ considering the lags of the residuals obtained after estimation of the both AR models (for II and PIS, respectively) for the first time window:** $m = 1$. The signal used in this case was from SUB, as in the main body of the manuscript.

Fig B presents the AR coefficients obtained for all of the discrete-time models $\mathbf{A}_m^{\{c\}}$, $m = 1, 2, ..., M$ considering the II and PIS conditions (Figs. BA and BB), respectively. The coefficients vary over time but are enclosed within upper and lower limits.

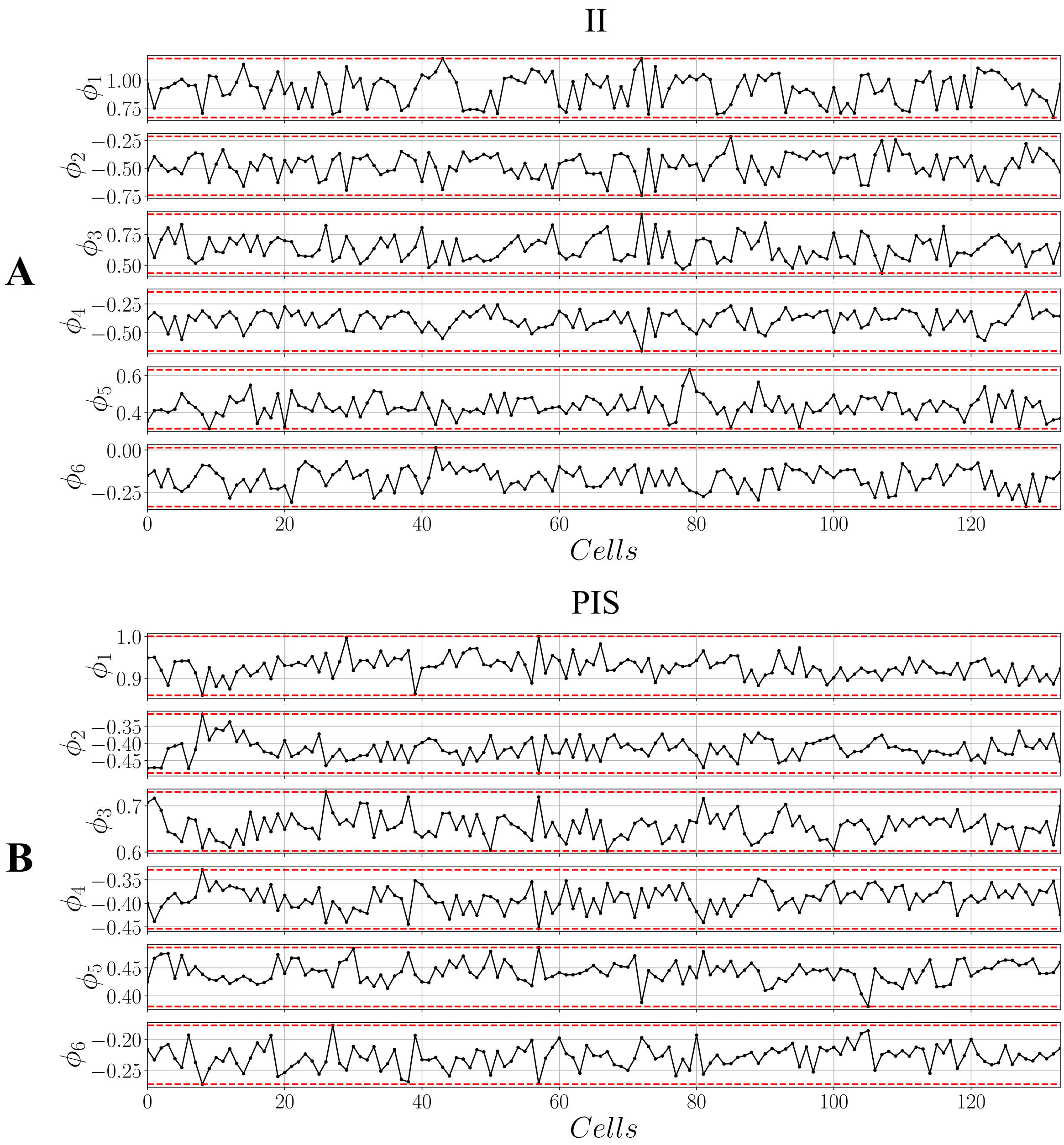


**Fig B. AR coefficients of the identified models for the II and PIS signals from SUB to design matrices $\mathbf{A}_m^{\{c\}}$, $m = 1, 2, .., M$ and the gains $\mathbf{L_{II}}$ and $\mathbf{G_{II}}$, or $\mathbf{L_{PIS}}$ and $\mathbf{G_{PIS}}$.** **–** represents the coefficients; **- -** represents their upper and lower limits.

Fig C presents the results for the performance of the controller when applied to signals coming from different hippocampal subfields. In general, it is efficient in attenuating the PIS activities while driving the system back to the II condition. Even though attenuation occurs, poorer results are seen for CA1, where a high fluctuation around the baseline occurs, possibly due to stronger background/noise activity.

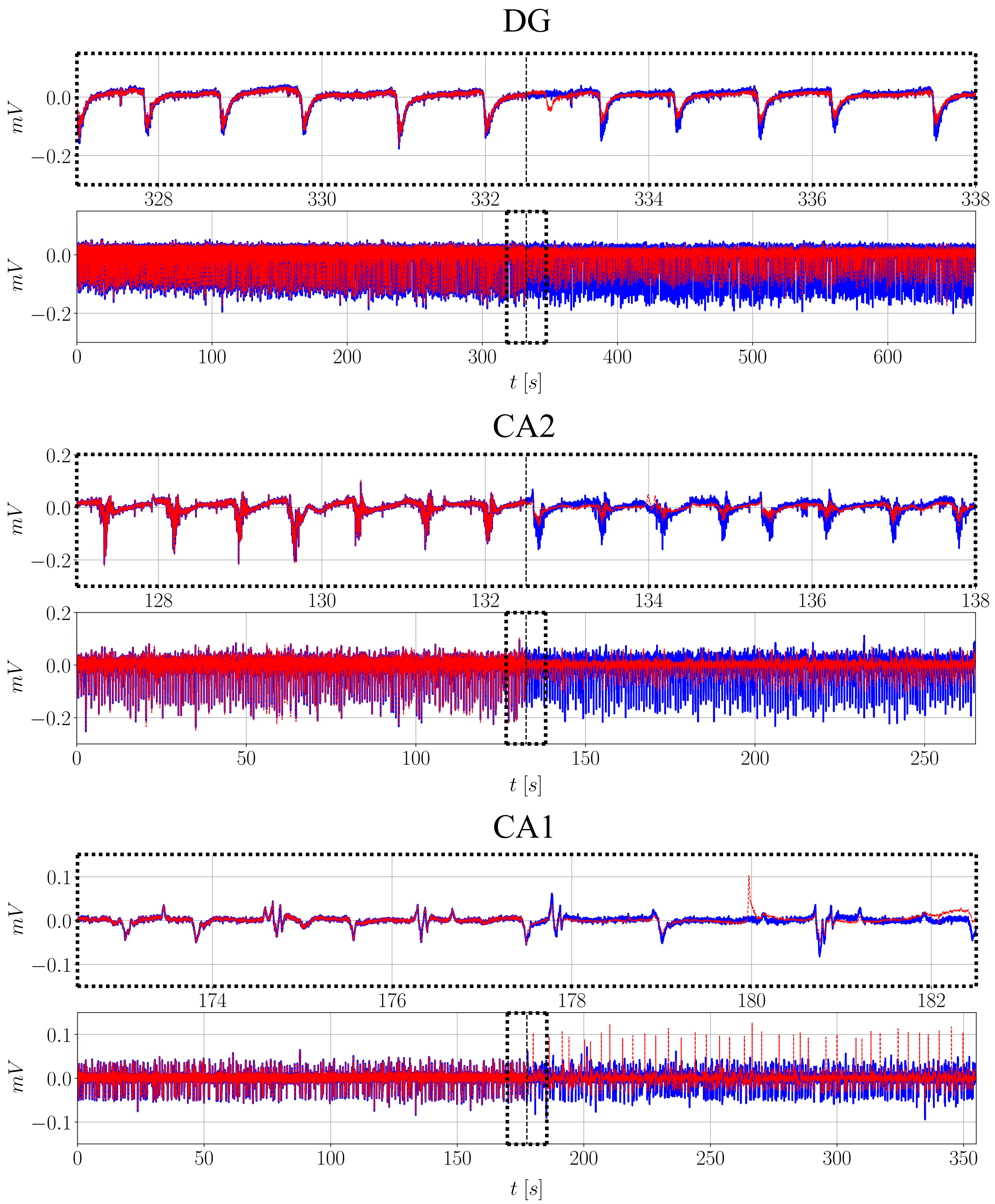


**Fig C. Performance of the controller when applied to signals coming from different hippocampal subfields: DG, CA2 and CA1**. – represents the uncontrolled activity; - - represents the controlled activity. In this case $\mathbf{L}_{\mathrm{PIS}}$ and $\mathbf{G}_{\mathrm{PIS}}$ are used without any constraints on the input. The observers and controllers were designed according to the available dataset, so results for CA3 and CA4 could not be generated due to the lack of PIS activity available for them.

Fig D shows the signal (number 21 from SUB) used to design both gains $\mathbf{L}_{\mathrm{II}}$ and $\mathbf{G}_{\mathrm{II}}$ in the hybrid approach covered in the main text, whereas $\mathbf{x}_{\mathrm{h}}$ (signal 18 from SUB) is the healthy reference signal to which drive the system. Note that the non-periodic feature is present in both 18 and 21, thus characterizing the II events. However, signal 21 has lower amplitude and was chosen for the design to avoid significantly higher gains. This suggests that amplitude is just as important as the spiking pattern when designing controllers and observers.

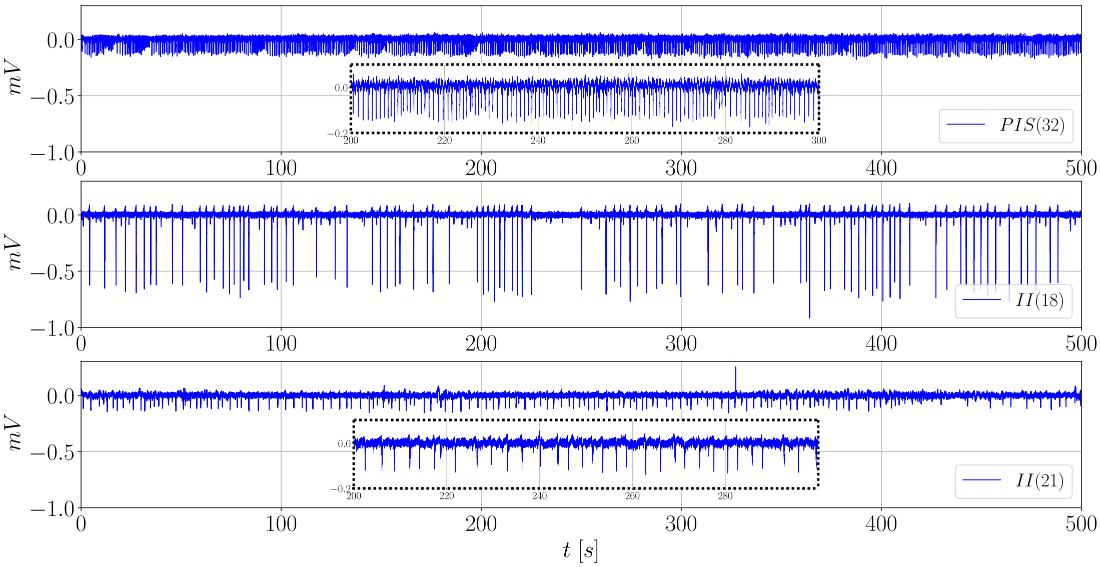


**Fig D. Signals used to design both gains $\mathbf{L_{II}}$ and $\mathbf{G_{II}}$ in the hybrid approach and the healthy reference $\mathbf{x_h}$.** In this case, signal number 21 from SUB is used for the design, whereas signal number 18 is used as a reference to which drive the system.
